# K_V_1.8 potassium channel subunits of vestibular hair cells affect mechanically-evoked hair cell and afferent signals and vestibulomotor function

**DOI:** 10.1101/2025.01.28.634388

**Authors:** Hannah R. Martin, Brandie Morris Verdone, Omar López-Ramírez, Merrill Green, Dana Silvian, Emily Scott, Kathleen E. Cullen, Ruth Anne Eatock

**Author notes:** Corresponding author:* **Author for Correspondence:** Ruth Anne Eatock, Ph.D. **Conflict of interest statement**: The authors declare no competing financial interests.

## Abstract

Vestibular hair cells (HCs) have large voltage-gated K^+^ conductances that shape the encoding and transmission of head motions and position. The K_V_1.8 (*Kcna10*) subunit is selectively expressed in amniote inner ear hair cells (HCs), and its deletion eliminates most of the depolarization-activated K^+^ conductance of type I and II HCs in the mouse utricle. Here we report effects of K_V_1.8 deletion on vestibular inner ear function and on vestibulomotor behaviors that stabilize posture, locomotion and gaze. In utricles excised from *Kcna10*^-/-^ and control mice, we deflected individual hair bundles in the extrastriolar sensory epithelium and recorded transduction currents and receptor potentials from HCs and nonquantal (NQ) postsynaptic potentials from calyceal afferent terminals on type I HCs. Based on these comparisons, K_V_1.8 speeds up receptor potentials in both HC types, increasing lowpass corner frequencies from ∼20 Hz to >50 Hz. K_V_1.8 also attenuates the receptor potential, but increases synaptic gain 5- to 10-fold, such that the postsynaptic calyx gain is little affected. Because *Kcna10* is not expressed in the central nervous system, these inner ear effects likely underlie deficits that we observed in *Kcna10*^-/-^ mice under vestibulomotor challenges: vestibulo-ocular reflex, head control, rearing activity, swimming posture, and narrow balance beam performance. We conclude that the effects of K_V_1.8-containing channels at the first vestibular synapse are significant for vestibulomotor function.

**Significance statement:** Voltage-gated potassium (K_V_) channels strongly affect the gain and dynamics of hair cell (HC) responses to current inputs. In mice, K_V_1.8 is the major K_V_ subunit in all vestibular HCs, and is not detected elsewhere in the nervous system. However, K_V_ 1.8’s influence on vestibulomotor behavior is not established. In this paper, we show that K_V_1.8 deletion changed receptor potential gain, speed, and displacement bandwidth, and reduced the efficacy of nonquantal transmission, the dominant mechanism by which type I HCs signal to postsynaptic calyceal terminals. Knockout mice were impaired on tasks that demanded unstable postures, lacked proprioceptive cues, or required precise timing, linking for the first time a single HC K_V_ channel to the fidelity of postural and gaze stabilization.

## Introduction

The vestibular inner ear of all vertebrates has a common macroscopic plan, reflecting the shared need to detect head motion and tilt. After stem amphibians moved onto land amniotes evolved and introduced not just a new auditory organ but also a new vestibular hair cell type (I) with a calyceal afferent synapse. The type I-calyx synapse uses nonquantal (NQ) transmission involving ion flow through low-voltage activated conductances in hair cell and calyx membranes facing each other across the synaptic cleft. The most prominent of these conductances is the type I HC’s large K^+^-selective conductance (g_K,L_ or g_KI_) that activates with depolarization over a very negative voltage range. In contrast, type II hair cells in the same epithelia are more conventional: they have A-type and delayed rectifier K^+^ conductances that activate positive to resting potential, and they transmit via glutamate-filled quanta (vesicles) to bouton afferent terminals. g_K,L_ plays a major role in changing the ionic composition of the synaptic cleft and enabling NQ transmission (Contini et al., 2017; Govindaraju et al., 2023). It has been natural to suggest that NQ transmission represents an adaptation to vestibular challenges encountered on land (Eatock, 2018). Indeed, NQ mechanisms transmit a faster and more faithful representation of the stimulus to the afferent spike initiation zone (Songer and Eatock, 2013; Govindaraju et al., 2023), features well-suited for driving swift and accurate vestibulomotor reflexes to compensate for rapid head motions.

We recently reported (Martin et al., 2024) that the major K^+^ conductances of type I and II HCs in the mouse utricle, despite their prominent physiological differences, all flow through ion channels that require K_V_1.8 subunits (gene name *Kcna10*). Whole-cell recordings in response to injected current showed that deleting K_V_1.8 produces large increases in HC membrane resistance in both HC types. Here we compare, for the extrastriolar zone of the mouse utricle, the effects of deleting K_V_1.8 on the gain and time course of mechanically-evoked receptor potentials from type I and II HCs and NQ postsynaptic potentials (PSPs) in calyces. To estimate the impact of K_V_1.8 on the type I-calyx synaptic transfer, we compare average type I HC and calyx responses to similar hair bundle stimuli.

We then ask how the prominent K_V_1.8 expression of vestibular hair cells affects vestibular system performance. In the original paper on these K_V_1.8-null mice, <u>Lee et al. (2013</u>) showed that vestibular HCs strongly express *Kcna10* and that *Kcna10*^-/-^ mice lack a vestibular-evoked potential (VsEP), a vibration-evoked electrical signal that can be recorded in anesthetized animals with scalp electrodes. The VsEP is very rapid (P1 latency ∼1 ms, Jones et al., 2011) and represents the synchronized spiking activity of afferents from the central (striolar) zone of the saccule (McCue and Guinan, 1994; Curthoys et al., 2016; Ono et al., 2020). The saccule is, like the utricle, an otolith organ sensitive to linear accelerations and head tilt, but its sensory epithelium is approximately vertical while the utricle’s is approximately horizontal. In many species, the saccule also detects moderately loud sounds. Given that *Kcna10* expression has not been detected in the brain (RNA, Lee et al., 2013) or in vestibular afferents (immunolabeling, Martin et al., 2024), its contribution to the VsEP presumably originates in vestibular HCs.

Here we describe the impact of K_V_1.8 deletion on other indicators of vestibular performance. Vestibular afferents that project to brainstem vestibular nuclei drive our fastest and most powerful reflexes: vestibulo-ocular reflexes (VOR) that maintain a stable visual field during head motions, vestibulocollic reflexes that stabilize head position, and vestibulospinal reflexes that correct limb and body position to stabilize posture and locomotion (Huterer and Cullen, 2002; Cullen, 2023). We compared VOR and other vestibulomotor behaviors between *Kcna10*^-/-^ mice and littermate controls to determine how the observed changes in peripheral signaling affect gaze stabilization and balance.

Together, our results illuminate how K_V_1.8 subunits condition receptor potentials and afferent signals in ways that enhance precise and fast compensation for head motions and posture. While the K_V_1.8-dependent subunit has the greatest effect in type I HCs, where it strongly reduces input resistance and contributes to NQ transmission to calyces, K_V_1.8-dependent subunits in type II HCs (g_A_ and g_DR_) also speed up type II receptor potentials and presumably their quantal transmission of glutamate to afferents.

## Materials and Methods

### Preparation

All procedures for handling animals followed the NIH Guide for the Care and Use of Laboratory Animals and were approved by the Institutional Animal Care and Use Committees of the University of Chicago (Animal Care and Use Protocol #72360) and Johns Hopkins University (Animal protocol #MO23M106). Procedures were performed at the University of Chicago unless otherwise stated. All mice belonged to a transgenic line with a knockout allele of *Kcna10* (referred to here as *Kcna10*^-/-^). Our breeding colony was established with a generous gift from Sherri Jones and Tom Friedman. As described more fully in Lee et al., 2013, the line was generated on a C57BL/6;129SvEv mixed background by replacing Exon 3 of the *Kcna10* gene with an IRES-bGeo/Purocassette. Mice in our colony were raised on a 12:12 light-dark cycle with access to food and water *ad libitum*.

Preparation, stimulation, and recording methods followed our previously described methods for the mouse utricle (Vollrath and Eatock, 2003). Chemicals were obtained from Sigma-Aldrich unless otherwise specified. Mice from a broad range of ages were used (hair cell physiology, P6-P60; calyx physiology, P14-P259; behavior, 1.3-8.4 months). Within these groups, specific questions may have been addressed with smaller age ranges; the median and range for each are provided in data tables. Animals were deeply anesthetized by isoflurane inhalation and decapitated. The head was bisected longitudinally and each half bathed in ice-cold, oxygenated Liebowitz-15 (L15) medium supplemented with 10 mM HEPES. The temporal bone was removed and the labyrinth was dissected to isolate the utricle and open the utricular sac. The utricular branch of the vestibular nerve was cut close to the utricular epithelium for HC recordings and after the vestibular ganglion for afferent recordings, allowing afferent arbors to remain attached to their cell bodies in the ganglion. The utricle was treated with proteinase XXIV (100 micrograms/mL, ∼10 mins, 22°C) to facilitate removal of the otoconia and attached gel layer, and mounted beneath two fine glass rods affixed at one end to a coverslip. The experiment chamber was mounted on a Zeiss Axio Examiner (A1) and viewed with Differential Interference Contrast (DIC) optics at 63X.

### Electrophysiology

We performed whole-cell recordings from hair cells and afferent calyceal terminals in the semi-intact epithelium with a patch clamp amplifier (HEKA Multiclamp EPC10) controlled by Patchmaster acquisition software. Output signals were filtered by the integrated HEKA Bessel filters (6-pole filter at 10 kHz plus 4-pole filter at 5 kHz), and sampled at 0.1 – 0.01 ms intervals. Recording electrodes were pulled (PC-100, Narishige) from soda lime glass (King’s Precision Glass R-6) and wrapped in paraffin to reduce pipette capacitance. Internal solution contained (in mM) 135 KCl, 0.5 MgCl_2_, 3 MgATP, 5 HEPES (2-[4-(2-hydroxyethyl)piperazin-1-yl]ethanesulfonic acid), 5 EGTA (ethylene glycol-bis(β-aminoethyl ether)-N,N,Nʹ,Nʹ-tetraacetic acid), 0.1 CaCl_2_, 0.1 Na-cAMP, 0.1 LiGTP, and 5 Na_2_CreatinePO_4_ and adjusted to pH 7.25 and ∼280 mmol/kg by adding ∼30 mM KOH, bringing total K^+^ to ∼165 mM. The external solution was L15 medium supplemented with 10 mM HEPES (pH 7.40, 310 ± 10 mmol/kg). Recordings were conducted at room temperature (22-25°C) to increase the lifespan of preparations and to avoid electrical noise and pipette drift that occurs with warming. Pipette capacitance and membrane capacitance transients were subtracted online with Patchmaster software. Series resistance (8-12 MΩ) was measured after rupture and compensated 0-80% with the amplifier such that mean remaining series resistance after online compensation was 9 ± 1 megaOhms (n=34 HCs). Potentials are corrected for uncompensated series resistance and liquid junction potential of ∼+4 mV, calculated with LJPCalc software (Marino et al., 2014).

### Transduction

We evoked MET currents and receptor potentials (V_RecP_) by displacing the hair bundle along its sensitive axis (Shotwell et al., 1981) with a stiff glass rod contacting the back of the hair bundle about halfway up the stereocilia staircase (**Fig. 1A**). The probe was pulled from borosilicate glass, tip width <1 micron, shank 16 mm, and fixed to a 1D piezoelectric bimorph element (5mm x 10 mm) (Corey and Hudspeth, 1979). Driving voltage to the bimorph was filtered with an 8-pole Bessel low-pass filter (Model 900, Frequency Devices, Ottawa, IL) at a corner frequency of 1 kHz. The probe’s time course was monitored offline by projecting the probe image onto the edge of a photodiode (PIN-6D, United Detector Technology, Hawthorne CA). The probe’s resonant frequency (2.25 kHz) was determined by averaging the first 4 periods of undamped oscillations evoked by an unfiltered step of driving voltage. The probe’s 10-90% rise time was ∼230 microseconds. At the end of each experiment, we used the experiment microscope and camera (QI-Click, QImaging) to calibrate the probe motion per driving voltage from the steady-state excursion of the probe tip during a sustained (1-s) step. For these experiments, the probe calibration was ∼400 nm/V and peak displacements were around 1.2 µm.

**Figure 1.**
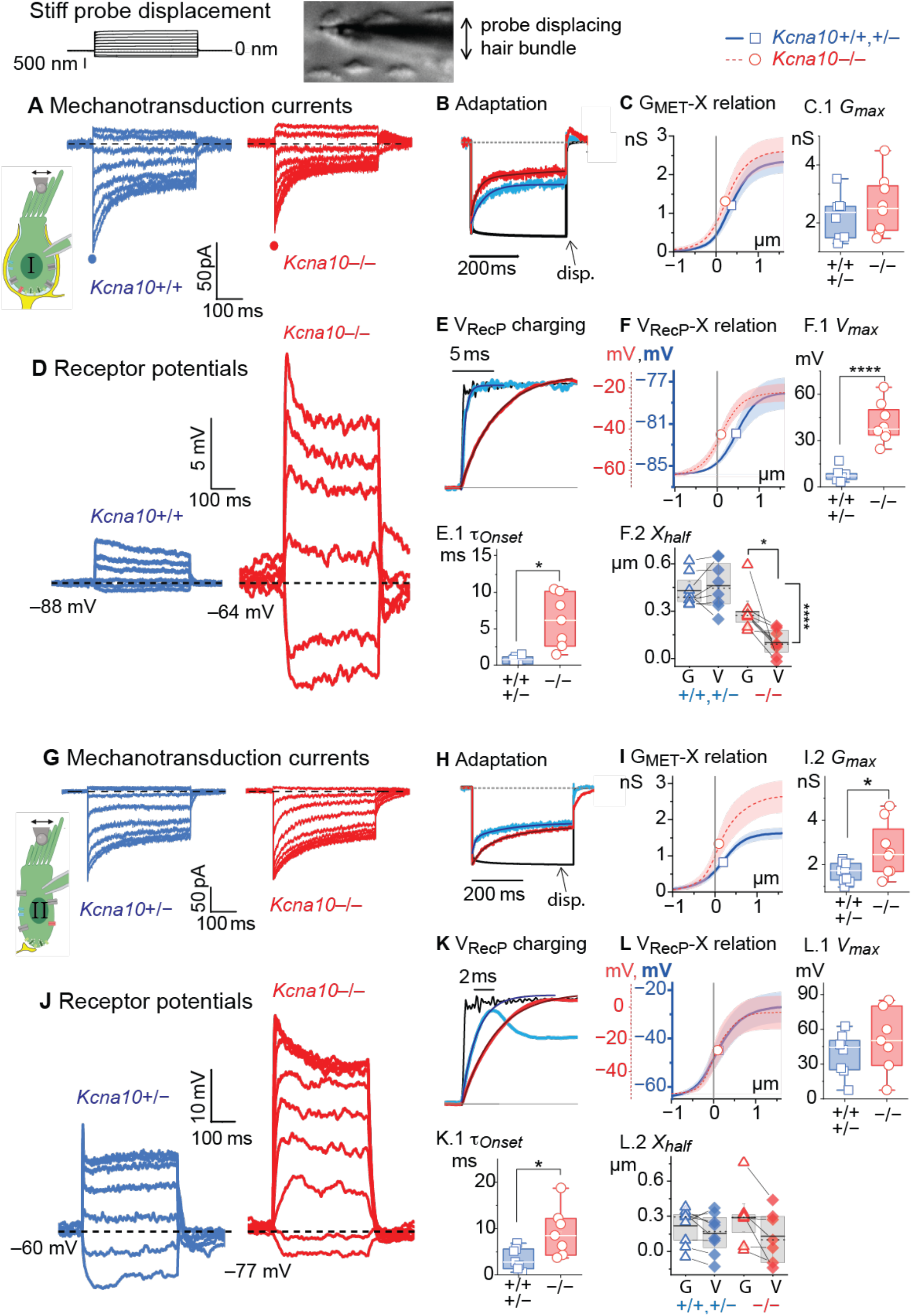
*Kcna10*^-/-^ hair cells have larger and slower receptor potentials. Top panel, Displacement stimulus (–0.3 to 1 µm) for all results. (**A**) Exemplar step-evoked MET currents from *Kcna10*^+/+^ at –87 mV, LES P17; *Kcna10*^-/-^ at –84 mV, MES P45. (**B**) Normalized current responses to +0.5 µm displacement; adaptation fit with a double exponential. Displacement (Disp) measured from offline photodiode recording (see Methods). (**C**) Mean +/- SEM. Fits of <u>Eq.1</u> and <u>2</u> to peak G-*X* relations from *Kcna10*^+/+,+/–^ (n=8) and *Kcna10*^-/-^ (n=8) type I HCs. *Open symbols*, *X_1/2_*. C.1, Boxplot of *G_Max_*. (**D**) Exemplar V_RecP_ from same cells as B. (**E**) Voltage responses overlaid with +0.25 µm displacement and exponential fit. E.1, *τ*_Onset_, responses to 30-470 nm displacements, Welch ANOVA p=0.008, g 2. (**F**) Mean ± SEM. Fits of <u>Eq.1</u> to peak V-X relations from *Kcna10*^+/+,+/–^ (n=8) and *Kcna10*^-/-^ (n=7). F.1, *V_Max_*, t-test p=6E-6, g 3.6. F.2, *X_half_*, differences in activation midpoint of each cell’s G-X and V-*X* relations. Top bar, p=0.03, *g* 1.4; side bar, p = 7E-5, *g* 3. (**G**) Type II HCs’ step-evoked I-MET obtained at –84 mV from *Kcna10^+/−^*, P20 LES; *Kcna10^-/-^*, P6 LES. (**H**) Normalized current responses to +0.5 µm displacement; adaptation fit with a double exponential. (**I**) Mean ± SEM. Fits of Eq. 2 to peak G-X relation from *Kcna10*^+/+,+/–^ (n=8) and *Kcna10*^-/-^ (n=8). *G_Max_*, ANOVA p=0.03, g 1.2. (**J**) Exemplar V_RecP_ in type II HCs. (**K**) Exemplar voltage responses to +0.25 µm displacements fit to Eq. 3. K.1, *τ*_Onset_, responses to 10-160 nm displacements, ANOVA p=0.04, g 1.2. (**L**) Peak V_RecP_ (X) averaged across cells. L.1, *V_Max_.* L.2, *X_half_*. Full statistics in Tables 1-2. Symbols: *p<0.05, **p<0.01, **** p<1E-5. Box plots: horizontal line, median; box, interquartile range; whiskers, furthest point.

**Table 1.**
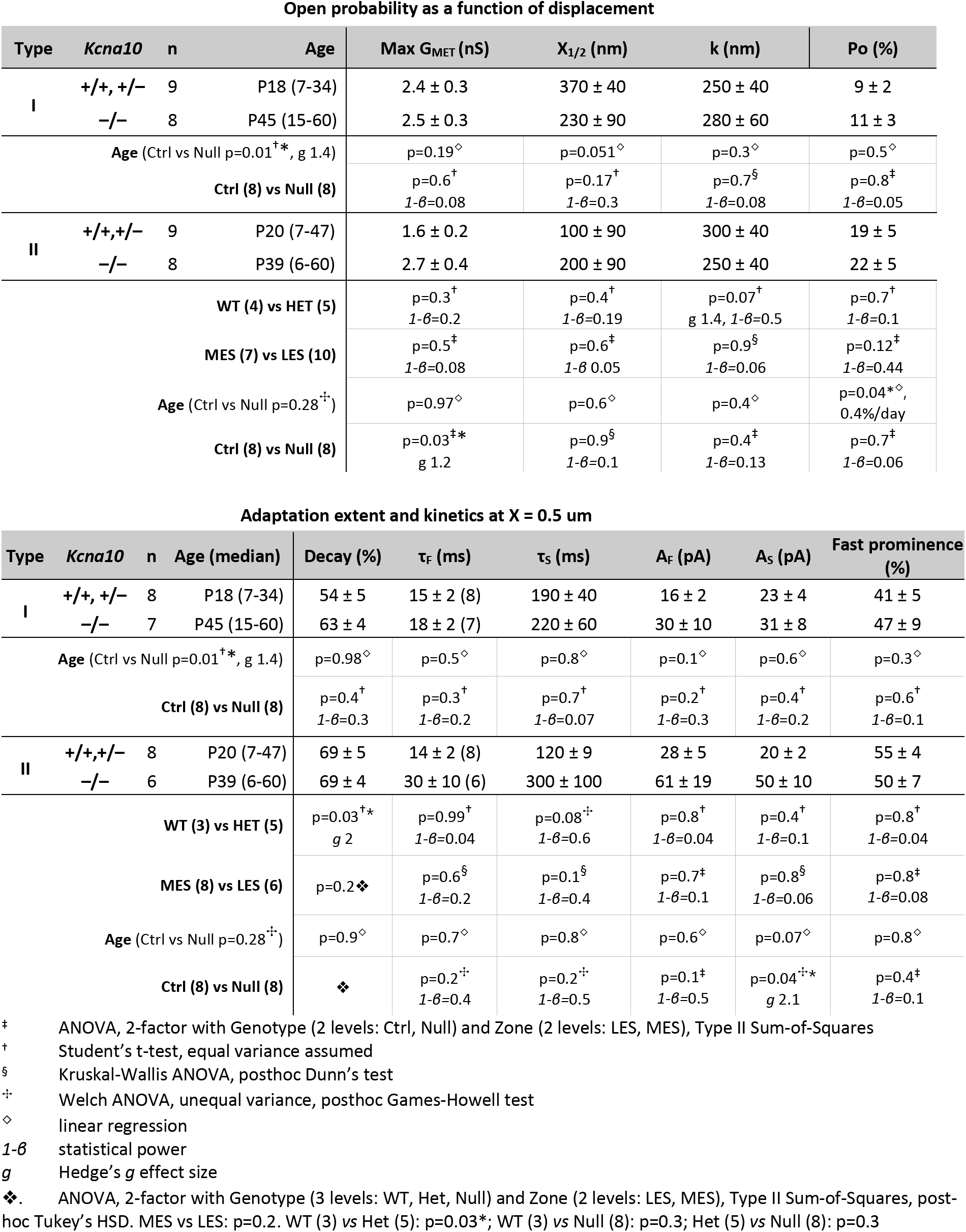
Step-evoked MET currents and adaptation in extrastriolar hair cells. Mean ± SEM (number of cells) Type II HC parameters were also compared across the two zones tested (lateral extrastriola, LES; medial extrastriola, MES) and between Kcna10+/+ (WT) and +/– (HET). Age is median postnatal day (range). X_1/2_ and k include X_1_ and k_1_, respectively, from fits to a second order Boltzmann function (Eq. 2).

| Open probability as a function of displacement |  |  |  |  |  |  |  |  |  |
|---|---|---|---|---|---|---|---|---|---|
| Type | *Kcna10* | n | Age | Max G<sub>MET</sub> (nS) | X<sub>1/2</sub> (nm) | k (nm) | Po (%) |  |  |
| I | +/+, +/− | 9 | P18 (7-34) | 2.4 ± 0.3 | 370 ± 40 | 250 ± 40 | 9 ± 2 |  |  |
|  | −/− | 8 | P45 (15-60) | 2.5 ± 0.3 | 230 ± 90 | 280 ± 60 | 11 ± 3 |  |  |
| **Age** (Ctrl vs Null p=0.01<sup>†</sup>\*, g 1.4) |  |  |  | p=0.19<sup>◇</sup> | p=0.051<sup>◇</sup> | p=0.3<sup>◇</sup> | p=0.5<sup>◇</sup> |  |  |
| **Ctrl (8) vs Null (8)** |  |  |  | p=0.6<sup>†</sup><br>*1-β*=0.08 | p=0.17<sup>†</sup><br>*1-β*=0.3 | p=0.7<sup>§</sup><br>*1-β*=0.08 | p=0.8<sup>‡</sup><br>*1-β*=0.05 |  |  |
| II | +/+,+/− | 9 | P20 (7-47) | 1.6 ± 0.2 | 100 ± 90 | 300 ± 40 | 19 ± 5 |  |  |
|  | −/− | 8 | P39 (6-60) | 2.7 ± 0.4 | 200 ± 90 | 250 ± 40 | 22 ± 5 |  |  |
| **WT (4) vs HET (5)** |  |  |  | p=0.3<sup>†</sup><br>*1-β*=0.2 | p=0.4<sup>†</sup><br>*1-β*=0.19 | p=0.07<sup>†</sup><br>g 1.4, *1-β*=0.5 | p=0.7<sup>†</sup><br>*1-β*=0.1 |  |  |
| **MES (7) vs LES (10)** |  |  |  | p=0.5<sup>‡</sup><br>*1-β*=0.08 | p=0.6<sup>‡</sup><br>*1-β* 0.05 | p=0.9<sup>§</sup><br>*1-β*=0.06 | p=0.12<sup>‡</sup><br>*1-β*=0.44 |  |  |
| **Age** (Ctrl vs Null p=0.28<sup>✢</sup>) |  |  |  | p=0.97<sup>◇</sup> | p=0.6<sup>◇</sup> | p=0.4<sup>◇</sup> | p=0.04*<sup>◇</sup>,<br>0.4%/day |  |  |
| **Ctrl (8) vs Null (8)** |  |  |  | p=0.03<sup>‡</sup>*<br>g 1.2 | p=0.9<sup>§</sup><br>*1-β*=0.1 | p=0.4<sup>‡</sup><br>*1-β*=0.13 | p=0.7<sup>‡</sup><br>*1-β*=0.06 |  |  |
| **Adaptation extent and kinetics at X = 0.5 um** |  |  |  |  |  |  |  |  |  |
| **Type** | ***Kcna10*** | **n** | **Age (median)** | **Decay (%)** | **τ<sub>F</sub> (ms)** | **τ<sub>S</sub> (ms)** | **A<sub>F</sub> (pA)** | **A<sub>S</sub> (pA)** | **Fast prominence (%)** |
| I | +/+, +/− | 8 | P18 (7-34) | 54 ± 5 | 15 ± 2 (8) | 190 ± 40 | 16 ± 2 | 23 ± 4 | 41 ± 5 |
|  | −/− | 7 | P45 (15-60) | 63 ± 4 | 18 ± 2 (7) | 220 ± 60 | 30 ± 10 | 31 ± 8 | 47 ± 9 |
| **Age** (Ctrl vs Null p=0.01<sup>†</sup>\*, g 1.4) |  |  |  | p=0.98<sup>◇</sup> | p=0.5<sup>◇</sup> | p=0.8<sup>◇</sup> | p=0.1<sup>◇</sup> | p=0.6<sup>◇</sup> | p=0.3<sup>◇</sup> |
| **Ctrl (8) vs Null (8)** |  |  |  | p=0.4<sup>†</sup><br>*1-β*=0.3 | p=0.3<sup>†</sup><br>*1-β*=0.2 | p=0.7<sup>†</sup><br>*1-β*=0.07 | p=0.2<sup>†</sup><br>*1-β*=0.3 | p=0.4<sup>†</sup><br>*1-β*=0.2 | p=0.6<sup>†</sup><br>*1-β*=0.1 |
| II | +/+,+/− | 8 | P20 (7-47) | 69 ± 5 | 14 ± 2 (8) | 120 ± 9 | 28 ± 5 | 20 ± 2 | 55 ± 4 |
|  | −/− | 6 | P39 (6-60) | 69 ± 4 | 30 ± 10 (6) | 300 ± 100 | 61 ± 19 | 50 ± 10 | 50 ± 7 |
| **WT (3) vs HET (5)** |  |  |  | p=0.03<sup>†</sup>\*<br>*g* 2 | p=0.99<sup>†</sup><br>*1-β*=0.04 | p=0.08<sup>✢</sup><br>*1-β*=0.6 | p=0.8<sup>†</sup><br>*1-β*=0.04 | p=0.4<sup>†</sup><br>*1-β*=0.1 | p=0.8<sup>†</sup><br>*1-β*=0.04 |
| **MES (8) vs LES (6)** |  |  |  | p=0.2❖ | p=0.6<sup>§</sup><br>*1-β*=0.2 | p=0.1<sup>§</sup><br>*1-β*=0.4 | p=0.7<sup>‡</sup><br>*1-β*=0.1 | p=0.8<sup>§</sup><br>*1-β*=0.06 | p=0.8<sup>‡</sup><br>*1-β*=0.08 |
| **Age** (Ctrl vs Null p=0.28<sup>✢</sup>) |  |  |  | p=0.9<sup>◇</sup> | p=0.7<sup>◇</sup> | p=0.8<sup>◇</sup> | p=0.6<sup>◇</sup> | p=0.07<sup>◇</sup> | p=0.8<sup>◇</sup> |
| **Ctrl (8) vs Null (8)** |  |  |  | ❖ | p=0.2<sup>✢</sup><br>*1-β*=0.4 | p=0.2<sup>✢</sup><br>*1-β*=0.5 | p=0.1<sup>‡</sup><br>*1-β*=0.5 | p=0.04<sup>✢</sup>\*<br>*g* 2.1 | p=0.4<sup>‡</sup><br>*1-β*=0.1 |

<sup>‡</sup> ANOVA, 2-factor with Genotype (2 levels: Ctrl, Null) and Zone (2 levels: LES, MES), Type II Sum-of-Squares
<sup>†</sup> Student's t-test, equal variance assumed
<sup>§</sup> Kruskal-Wallis ANOVA, posthoc Dunn's test
<sup>✢</sup> Welch ANOVA, unequal variance, posthoc Games-Howell test
<sup>◇</sup> linear regression
*1-β* statistical power
*g* Hedge's *g* effect size
❖. ANOVA, 2-factor with Genotype (3 levels: WT, Het, Null) and Zone (2 levels: LES, MES), Type II Sum-of-Squares, post-hoc Tukey's HSD. MES vs LES: p=0.2. WT (3) *vs* Het (5): p=0.03\*; WT (3) *vs* Null (8): p=0.3; Het (5) *vs* Null (8): p=0.3

**Table 2.**
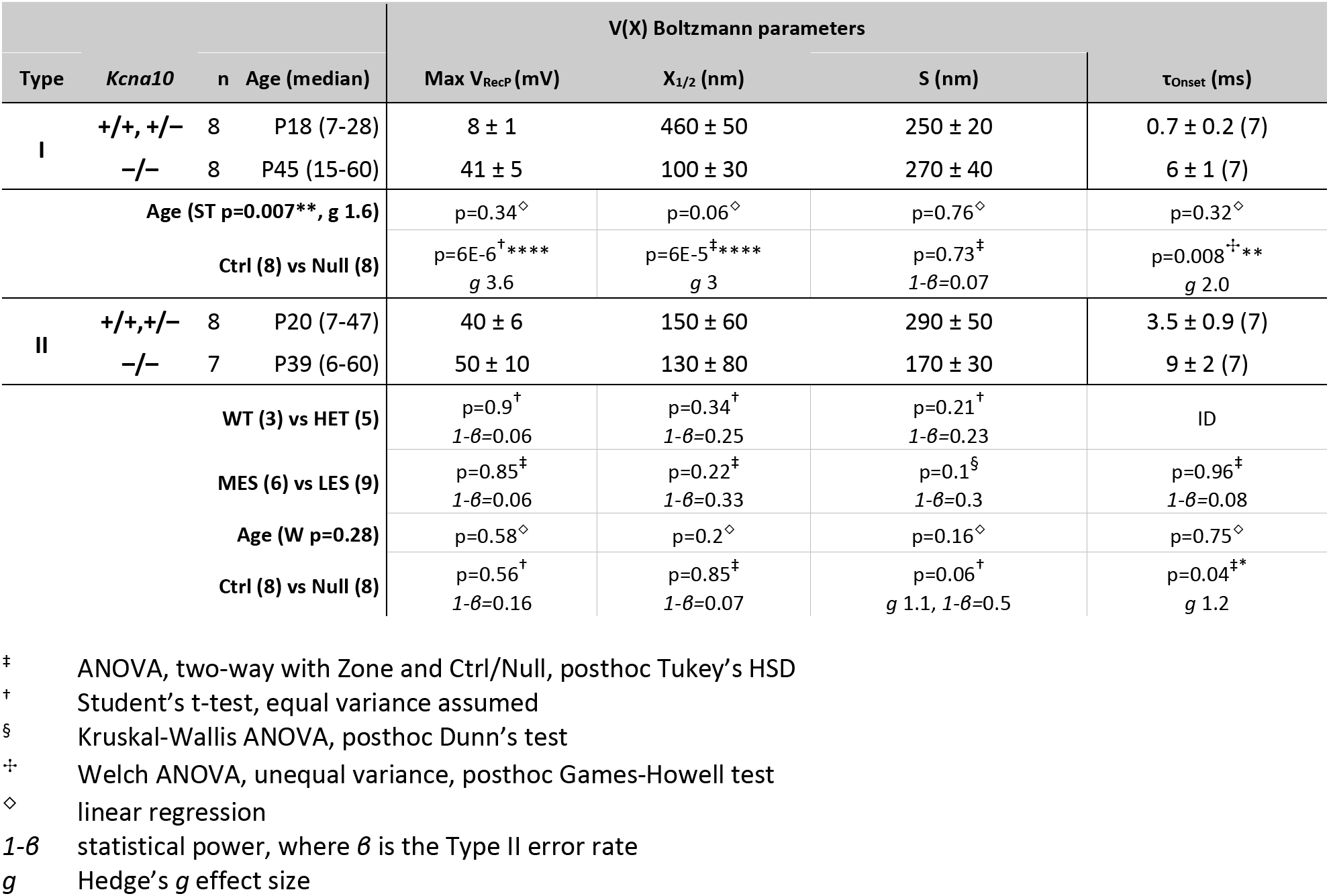
Step-evoked receptor potentials in extrastriolar hair cells. Mean ± SEM (number of cells). Type I HC parameters were compared by Age and Genotype. Type II HC parameters were also compared across the two zones tested (lateral extrastriola, LES; medial extrastriola, MES) and between *Kcna10+/+* (WT) and *+/–* (HET). Age is median postnatal day (range).

|  |  |  |  | **V(X) Boltzmann parameters** |  |  |  |
|---|---|---|---|---|---|---|---|
| **Type** | ***Kcna10*** | **n** | **Age (median)** | **Max V<sub>RecP</sub> (mV)** | **X<sub>1/2</sub> (nm)** | **S (nm)** | **τ<sub>Onset</sub> (ms)** |
| **I** | **+/+, +/−** | 8 | P18 (7-28) | 8 ± 1 | 460 ± 50 | 250 ± 20 | 0.7 ± 0.2 (7) |
|  | **−/−** | 8 | P45 (15-60) | 41 ± 5 | 100 ± 30 | 270 ± 40 | 6 ± 1 (7) |
| **Age (ST p=0.007\*\*, g 1.6)** |  |  |  | p=0.34<sup>◇</sup> | p=0.06<sup>◇</sup> | p=0.76<sup>◇</sup> | p=0.32<sup>◇</sup> |
| **Ctrl (8) vs Null (8)** |  |  |  | p=6E-6<sup>†</sup>\*\*\*\*<br>*g* 3.6 | p=6E-5<sup>‡</sup>\*\*\*\*<br>*g* 3 | p=0.73<sup>‡</sup><br>*1-β*=0.07 | p=0.008<sup>✢</sup>\*\*<br>*g* 2.0 |
| **II** | **+/+,+/−** | 8 | P20 (7-47) | 40 ± 6 | 150 ± 60 | 290 ± 50 | 3.5 ± 0.9 (7) |
|  | **−/−** | 7 | P39 (6-60) | 50 ± 10 | 130 ± 80 | 170 ± 30 | 9 ± 2 (7) |
| **WT (3) vs HET (5)** |  |  |  | p=0.9<sup>†</sup><br>*1-β*=0.06 | p=0.34<sup>†</sup><br>*1-β*=0.25 | p=0.21<sup>†</sup><br>*1-β*=0.23 | ID |
| **MES (6) vs LES (9)** |  |  |  | p=0.85<sup>‡</sup><br>*1-β*=0.06 | p=0.22<sup>‡</sup><br>*1-β*=0.33 | p=0.1<sup>§</sup><br>*1-β*=0.3 | p=0.96<sup>‡</sup><br>*1-β*=0.08 |
| **Age (W p=0.28)** |  |  |  | p=0.58<sup>◇</sup> | p=0.2<sup>◇</sup> | p=0.16<sup>◇</sup> | p=0.75<sup>◇</sup> |
| **Ctrl (8) vs Null (8)** |  |  |  | p=0.56<sup>†</sup><br>*1-β*=0.16 | p=0.85<sup>‡</sup><br>*1-β*=0.07 | p=0.06<sup>†</sup><br>*g* 1.1, *1-β*=0.5 | p=0.04<sup>‡\*</sup><br>*g* 1.2 |

<sup>‡</sup> ANOVA, two-way with Zone and Ctrl/Null, posthoc Tukey’s HSD
<sup>†</sup> Student’s t-test, equal variance assumed
<sup>§</sup> Kruskal-Wallis ANOVA, posthoc Dunn’s test
<sup>✢</sup> Welch ANOVA, unequal variance, posthoc Games-Howell test
<sup>◇</sup> linear regression
*1-β* statistical power, where *β* is the Type II error rate
*g* Hedge’s *g* effect size

The resting probe position evoked 16 ± 2 % (n = 38) of maximum MET currents [percent P_o_ = 100*(I_X=0_ – I_min_)/(I_max_ – I_min_)], consistent with our previous estimates on rodent vestibular HCs with similar methods (Vollrath and Eatock 2003; Songer and Eatock 2013). Holding potential ranged from –92 to –74 mV (median –84 mV). Holding potentials below –84 mV were used for some control type I HCs to fully deactivate g_K,L_. If activated, g_K,L_ can reduce voltage clamp quality by reducing the ratio of *R*_in_ to *R*_S_, as illustrated in **Supplemental Figure 1** through a model of an equivalent voltage circuit (Lei et al., 2020).

### Transduction Analysis

For data analysis, fitting, and plotting, we used OriginLab software (Northampton, MA) and custom MATLAB scripts.

#### Fitting sensitivity and time course of transduction currents

*G-X curves*. For each step, transduction current was taken as the maximum absolute change from baseline (I_Peak_). We then subtracted the I_Peak_ at the most inhibitory displacement, where I_MET_ open probability was lowest, and converted the resulting values to conductance (G) assuming a reversal potential of +2.6 mV (Kros et al., 1992, for mouse cochlear hair cells in similar solutions). The conductance values were then fit with a first-order Boltzmann equation (Eq. 1) and a second-order equation (Eq. 2), with the better fit selected on the basis of adjusted R^2^ and visual inspection. Eq. 2 was applied to ∼30% (3/9) of control type I HCs, ∼70% (5/7) of *Kcna10*^-/-^ type I HCs, 100% (8/8) of control type II HCs, and ∼60% (5/8) of *Kcna10*^-/-^ type II HCs. For each term, *G_min_* and *G_max_* are the saturating minimum and maximum G values of the total curve. *X_1_, X_2_ and X_half_* values are the midpoints, and *k* values are slope factors. In **Table 1**, cells fit by Eq. 2 are represented by *X_1_* and *k_1_*. V_RecP_-X curves were all fit by Eq. 1.

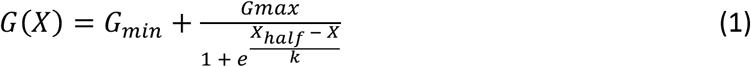

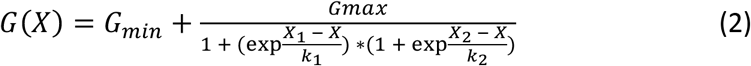

#### Adaptation time course

Adaptation of transduction currents for a displacement step to *X_half_* was well fit with Eq. 3. In 28/29 cells, 2 adapting terms provided a significantly better fit than one. *I_SS_* is steady-state current, *A_f_* and *A_s_* are amplitudes of fast and slow adaptation, and *τ_f_* and *τ_s_* are time constants of fast and slow adaptation. The cell well-fit by a single adapting term was a *Kcna10*^+/+^ type II from the MES at P49, with adaptation time course and amplitude (*τ* = 60 ms, *A* = 56 pA) intermediate between the fast and slow terms for the other cells.

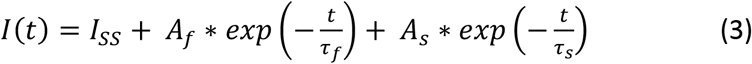

#### Frequency tuning

To capture frequency tuning, we recorded HC transduction currents or receptor potentials or calyx postsynaptic potentials to a bundle displacement protocol comprising eight 5-cycle sinusoidal bursts at frequencies incremented from 2 to 100 Hz (Songer and Eatock, 2013). We estimated the AC gain and phase by fitting each response burst with sine waves and referencing the fit to the corresponding bundle displacement, producing gain (pA/μm or mV/μm) and phase (degrees) relative to the stimulus. For each cell, phase- and gain-frequency curves were simultaneously fit with transfer functions (Equations 4a-b and 5a-d) to arrive at a single, shared corner frequency. These transfer functions have a characteristic order that sets the roll-off steepness and phase shift. We empirically determined the best order (*α*, *β*) for each group of cells or measurements: the high-pass component of transduction currents (*α* = 0.398), the high-pass component of receptor potentials (*α* = 0.338), and the lowpass component of receptor potentials and PSPs (*β* = 0.687). In each case, the best order was <1, indicating less gain roll-off than a single-pole filter (gain 6 dB/octave, and phase shift 90°).

For transduction currents, we fit gain-frequency curves to Eq. 4a, where A is the fitted maximal gain. High-pass (adaptation) fits to transduction current gain were restricted to 2–20 Hz. The corner frequency produced by transducer adaptation falls in this range, as expected from the measured adaptation time constants of 15–30 ms (Table 1). Roll-off was ∼3 dB/octave, consistent with adaptation distributed over multiple time constants.

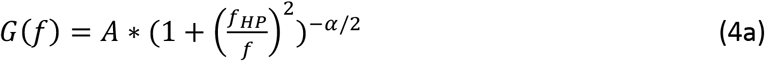

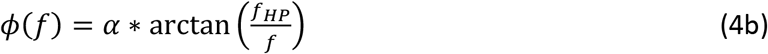

For receptor potentials and PSPs, gain-frequency and phase-frequency curves were fit by the lowpass and bandpass transfer functions described by Equations 5a-d. Gain-frequency curves from all PSPs and most receptor potentials were fit with Eq. 5a. However, in 50% (4/8) of control type II HC receptor potentials, the gain-frequency curve required instead fitting with a bandpass transfer function (Eq. 5b) because the curve had a noticeable high-pass (adaptation) component.

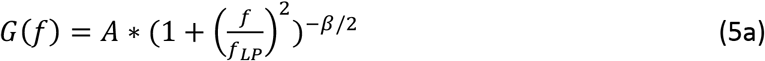

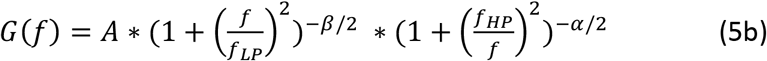

Receptor potential phase-frequency curves required fitting with a bandpass transfer function (Eq. 5c) to account for their phase lead at low frequencies. For Eq. 5b and 5c, *f_HP_* was empirically determined to be 5.36 Hz for all receptor potentials. PSP phase-frequency curves did not have a phase lead and were fit with only the lowpass function, Eq. 5d.

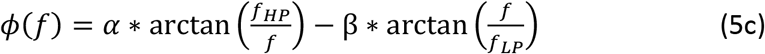

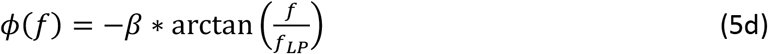

#### Synaptic transfer function

Synaptic gain was calculated as the ratio of the average peak displacement-evoked potentials in calyces and type I HCs, or *V*^-^*_Calyx_*/*V*^-^*_HC_*. We used the following methods to estimate the standard errors (*σ*_x-_) of the transformed parameters in **Figure 4D** and Table 4. We used (Equation 6) the delta method (Oehlert, 1992) to estimate the *σ_v/v_* of synaptic gain from the means (*V*^-^*Calyx*, *V*^-^*HC*) and their standard errors (*σ_V_*B,*Calyx, σ_V_*B,*HC*).The covariance terms *σ_calyx,HC_, σ_HC,calyx_* were allowed to be 0 because the calyx and type I HC recordings were obtained from different preparations, although we note that calyx potentials depend on HC potentials.

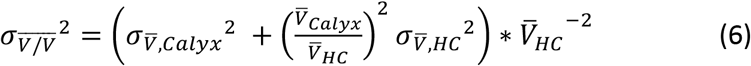

Synaptic phase (*φ*_DEF_) was the difference between the population averages of AC phase re: stimulus (bundle deflection), or *φ*^‒^*_Calyx_* − *φ*^‒^*_HC_*. *σ* of synaptic phase was the root sum of squares of *φ*^‒^*_Calyx_* and *φ*^‒^*_HC_*.

### Behavior

Animals were handled and introduced to the behavior room ≥24 hours before experiments.

#### Balance Beam

Animals were allowed 5-10 minutes to acclimate to the testing room. Mice were trained for 3 trials on a wide beam (12 mm wide), rested for 10 min, and then tested for 3 trials on a narrow beam (6 mm wide). To incentivize mice to cross the flat beam, we placed at the starting position a bright lamp and at the end of the beam a dark goal box (5 in x 5 in x 5 in) filled with food and bedding from their home cage. Mice were given at least 20 s to rest in the end box between trials. The beam was cleaned with Clidox-S between mice.

Videos of the trial were at 30 frames per second (fps) with a lateral view of the beam. Behavior in the videos was scored manually by two investigators blinded to genotype. The crossing time was recorded as the time between the nose crossing the starting line and the nose crossing the finish line (64 cm). Crossing time included non-ambulatory epochs, including while the mouse was being rescued and while the mouse stopped and sniffed the air. Inspection of the control and null animal performances produced several additional candidate behaviors for scoring: “beam slip”, hanging upside down on the beam and needing rescue to finish (rescue time was included in total time to cross); “tail wrap”, wrapping the tail around the beam instead of having it in the air; and “paw slip”, a single paw slipping off the beam without the mouse falling off the beam.

#### Swim test

A clear aquarium (38 cm long x 30 cm high x 25 cm wide) was filled with warm water (30°C) and placed in front of a white background. Animals were placed directly into the water one at a time. Videos (30 fps) of the swim were recorded from the side. After 30 seconds, animals were retrieved, dried with paper towels, and placed in a cardboard box under a heat lamp. For analysis, videos trimmed to the sections where the animal was swimming straight along the aquarium side closest to the camera. Two genotype-blind scorers marked body parts and calculated body angles in FIJI (Enthoven et al., 2004) in eight frames where the animal is swimming parallel to the camera view. This number of frames was empirically set after exploring the variability of the results, and a random number generator was used to pick frames for each scorer (repeat frames were not allowed).

#### Open field

Animals were placed in open field boxes (ENV-510, 27 x 27 x 20.3 cm); lighting was dim (16 lux) to reduce stress. After 10 minutes of acclimation, mouse movement was recorded for 1 h via infrared beam interruption. Activity Monitor 7 software (Med Associates, Inc, Fairfax, VT) returned data on the location of the mouse through time, and calculated Distance Travelled, Average Ambulatory Speed, Rearing Event Count, and Rearing Time. Centrophobism was calculated as percent of time spent outside the 8 cm x 8 cm central square.

#### Rotarod

We used a rotarod (Columbus Instruments) with large dowels (diameter = 7 cm). We used a 2-day protocol, one for training and one for testing, based on other studies of vestibular function (Tung et al., 2014; Jones et al., 2018; Rahman et al., 2022). On Day 1, there were 6-8 training trials: mice were initiated at 5 rpm then accelerated to 44 rpm at a rate of 2.4 rpm per 4 seconds, with 30-s rest between trials. On Day 2, the same protocol was used with 4 refresher trials, a 20-minute break, and then 3 test trials.

#### DigiGait

Mice were placed on a treadmill-like apparatus with a clear belt and an internal camera mounted below the mouse as it runs on the belt. Mice were recorded in slow (15 cm/s) and fast (25 cm/s) trials. For analysis, body parts were labelled for 35-70 frames per mouse for both the slow and fast trials. These labelled frames were used to train a DeepLabCut model in a Google Colab environment, which then applied labels to all frames. We analyzed gait parameters from these body part positions using custom Matlab functions and published equations (Fiker et al., 2020).

#### Grip Strength

Mice were allowed to grasp an accessory grasping grid with all four paws while an experimenter pulled back on their tail. The grip strength meter (BioSeb) reported the maximum pull-back force reached before the mouse lost its grip on the grid.

### Vestibulo-ocular reflex

Vestibulo-ocular reflex (VOR) testing was performed at Johns Hopkins University. Mice were head-restrained using a surgically implanted custom headpost, as previously described (Beraneck and Cullen, 2007). The mice were secured to a motion platform (hexapod: BREVA Dynamique - SN 21135, SYMETRIE) surrounded by a visual environment consisting of black and white stripes with a visual angle width of 5°, as described (Beraneck and Cullen, 2007). Eye positional movement data were sampled at 200 Hz with a platform-mounted camera (Firefly S USB3, FLIR) positioned ∼10 mm perpendicular to the mouse’s eye. We used custom pupil-tracking code to determine the horizontal and vertical components of 3D eye positional movement. The velocity of the platform (head velocity) was sampled at 500 Hz with a MEMs inertial measurement unit (IMU; ICM-46288-P, InvenSense/TDK) mounted on the headpost.

To evoke a VOR response, the motion platform delivered rotational sinewaves at 0.4, 0.6, 0.8, 1, 2, 5, 10, 15, and 20 Hz with peak velocity of ±15 deg/s and in both light and dark conditions. Synchronization pulses from IMU samples and camera frames were collected using National Instruments systems (PXI-6133) and Spike GLX software at a sampling rate of 20 kHz. These pulses were used to synchronize the eye and head movement data, which were then resampled at 1 kHz for analysis in MATLAB (2022b, Mathworks).

To compute the gain and phase of the VOR responses, eye position signals were differentiated to obtain eye velocity data. Segments of the eye movement data containing saccades or quick phase movements were identified and excluded from the analysis. Eye and head signals were low-pass filtered with a fourth-order Butterworth filter with a corner frequency set at twice the stimulation frequency plus one (Hz). A least-squares optimization was performed to determine the gain and phase at each frequency (see Beraneck and Cullen, 2007 for details). The estimated VOR gain and phase were then plotted as the mean ± SEM for each group.

### Instrumented Behavioral Assessments

Also at Johns Hopkins University, we assessed head motions during open field, swim, and balance beam tests. The tasks were similar to those described above, but were performed at the Johns Hopkins site where we had the capacity to affix a miniature IMU to each mouse’s head to quantify its angular rate and linear acceleration, as previously described (Beraneck et al., 2008; Chang et al., 2022). For the open field and swim assessments, head motions were recorded while mice performed the task in three 1-minute trials with 1 minute intertrial intervals. For balance beam, mice crossed a narrow (6-mm wide) beam for 3 trials with 20 seconds inter-trial intervals, now with an IMU affixed to the head. Head orientation during swim was calculated from IMU data as previously described (Hughes et al., 2024).

### Experimental Design and Statistical Analyses

Data normality was assessed with the Shapiro-Wilk test for n≤50 and the Kolmogorov-Smirnov test for other sets. We assessed homogeneity of variance with Levene’s test and, when variance was non-homogeneous, used a Welch ANOVA with the posthoc Games-Howell test. For data that were not normally distributed, we used the non-parametric Kruskal-Wallis ANOVA (KWA) with posthoc Dunn’s test.

With normal data and homogeneous variance, we used ANOVA with posthoc Tukey’s test. ANOVAs were Type I for animal behavior because cohorts were well-balanced and there was a reasonable chance of interaction between weight, sex, age, and genotype. ANOVAs were Type II for electrophysiology experiments that involved bundle displacement because these had small sample sizes (5-13 cells per group), unbalanced cohorts, and low expected chance of interaction between factors such as sex and age. Sample sex distribution is found in **Table S6**. To screen for any sex differences in transduction and receptor potential data, we ran a second series of statistical tests on combined type I and type II hair cell datasets that had greater power (**Table S7**).

We compared gain- and phase-frequency curves between groups of cells by a mixed one-way ANOVA: the within-factor was frequency, the between-factor was genotype, and an interaction term was included. If the interaction term was significant, then posthoc t-tests were performed at each frequency without a correction for multiple comparisons because measurements across frequencies are physiologically correlated and not independent.

Effect size is measured as Hedge’s g (*g*), as appropriate for our small samples. Our studies covered a wide range of animal age. Some cohorts have different median ages but overlapping age ranges. Behavioral tests and hair cell tests are largely non-overlapping with each other, with behavioral tests done past the age of sexual maturity (P40), and hair cell tests mostly below that age. To assess age dependence, we used linear models that included covariates of genotype and zone.

VOR gain and phase values were compared using two-way repeated measures ANOVA with post hoc Bonferroni corrections. For power spectra analysis, an independent sample permutation test was used to test significant differences between two groups (see <u>Hughes et al., 2024</u> for swim assessment details and (Ono et al., 2026) for details on power spectra analyses).

## Results

To explore the role of K_V_1.8 in transmission of head-motion signals, we first recorded mechanically-stimulated responses from type I and II HCs and calyces from the medial and lateral extrastriolar zones (MES, n=13 HCs; LES, n=19 HCs, n=11 calyces) of the utricular epithelium of mice (n=45) between postnatal day (P) 7 and P200. Hair bundles were displaced by a stiff probe with step and sinusoidal waveforms. We compared mechanotransduction (MET) currents and receptor potentials (V_RecP_) in type I and II HCs from homozygous knockout (*Kcna10*^-/-^) animals and their wildtype (*Kcna10*^+/+^) or heterozygote (*Kcna10*^+/–^) littermates. We pooled *Kcna10*^+/+^ and *Kcna10*^+/–^ data as “control” when they did not differ significantly.

To assess whether upstream changes in receptor potential waveform affected postsynaptic signaling, we next compared postsynaptic potentials (PSPs) in calyces of *Kcna10*^+/–^ (control) and *Kcna10*^-/-^ animals. As for the HC recordings, stimulation was directly applied to individual type I HBs providing input to the recorded calyces.

Finally, we compared the performance of *Kcna10*^-/-^ and control mice for several vestibulomotor activities: VOR, open field arena, balance beam, swimming, rotarod, and gait.

## I. Mechanically evoked signals in hair cells and calyces

### K_V_1.8 channels reduce receptor potential gain and latency in both hair cell types

Transduction was stimulated by pushing a calibrated stiff probe against the back of a hair bundle in its preferred direction. When step displacements open mechanotransduction (MET) channels at the tips of the hair bundles, inward MET currents (I_MET_) flow and subsequently adapt, or desensitize, by up to 70% (**Fig. 1A**). In both control and *Kcna10*^-/-^ type I HCs, I_MET_ declined (adapted) with similar kinetics described by a double exponential (**Fig. 1B**). I_MET_ was converted to conductance (G) to control for differences in holding potential across cells, and G_MET_(X) relations had an activation midpoint (X_1/2_) ∼300 nm (**Table 1**, Eq. 1, **Fig. 1C**). Genotype differences were not significant in maximal G_MET_, X_1/2_, displacement sensitivity (slope, S), or resting open probability (P_O_).

Absence of K_V_1.8 strongly altered the magnitude and time course of receptor potentials, consistent with the involvement of Kv1.8 subunits in the dominant K_V_ conductances of both HC types: g_K,L_ in type I and both g_A_ and g_DR_ in type II (Martin et al., 2024). Resting membrane potential (V_rest_) was more hyperpolarized in control than *Kcna10*^-/-^ type I HCs (**Fig. 1D**) because g_K,L_ (missing in the mutants) drives V_rest_ closer to E_K_. Control type I HCs produced small, fast receptor potentials that closely resembled the displacement step, because open g_K,L_ channels reduce input resistance (R_In_) at V_Rest_, shunting the potential (*V* = *IR*, **Fig. 1D**) and thus minimizing changes in the activation of voltage-gated channels during the step stimulus. In contrast, *Kcna10*^-/-^ type I HCs produced receptor potentials that were initially large, reflecting high R_In_, but decreased as residual (non-K_V_1.8) K_V_ conductances opened in response to depolarization (**Fig. 1D**).

Differences in R_In_ strongly affected V_RecP_ charging time (membrane time constant, *τ_m_* = *R_In_C_m_* (**Fig. 1E**). In control type I HCs, V_RecP_ occurred as fast as the stimulus, while high *τ_m_* slowed V_RecP_ in *Kcna10*^-/-^ type I HCs.

Receptor potential-displacement curves (**Fig. 1F**) showed that, relative to controls, *Kcna10*^-/-^ type I HCs have X_1/2_ shifted by ∼–300 nm. We further analyzed this shift by comparing X_1/2_ values from G_MET_ and V_RecP_ relations obtained from the same cell. In *Kcna10*^-/-^ type I HCs, X_1/2_ was shifted by ∼–100 nm for V_RecP_ relative to G_MET_ (p=0.03, paired sample t-test, n=6 cells, *g 1.4*), with no significant shifts observed in control HCs. *Kcna10*^-/-^ type I HCs alone featured this negative X_1/2_ shift because their larger V_RecP_ spans the activation range of the residual (non-K_V_1.8) g_DR_; once activated, g_DR_ resists further depolarization and compresses the V_RecP_-X curve such that the apparent X_1/2_ is more negative. Since g_K,L_ reduces the size of V_RecP_, control type I HCs don’t exhibit V_RecP_-X compression.

The properties of extrastriolar type II HC MET currents (**Fig 1G-I**) were generally similar to those of type I HCs. In both *Kcna10*^-/-^ and control type II HCs, I_MET_ also activated rapidly and declined with similar kinetics described by a double exponential (**Fig 1H**). However, G_MET_ was notably greater in *Kcna10*^-/-^ type II than control type II HCs (**Fig 1I**), an effect not due to unbalanced cohorts as the *Kcna10*^-/-^ and control type II HC datasets did not differ significantly by age or zone (Table 1).

Control type II HCs produced receptor potentials with an initial sharp positive-going peak that relaxed as g_A_’s rapid activation and a decaying driving force repolarized the membrane towards equilibrium (**Fig. 1J**) (Hudspeth and Lewis, 1988). *Kcna10*^-/-^ type II HCs produced receptor potentials that were larger because of higher input resistance and relaxed more slowly, reflecting the absence of Kv1.8-dependent g_A_ and g_DR_.

In **Figure 1K**, control type II HCs’ V_RecP_ had shorter rise times because of their lower input resistance, and repolarized more quickly because g_A_ has faster activation kinetics than the residual delayed rectifier channels in *Kcna10*^-/-^ type II HCs (Martin et al., 2024). Comparing *Kcna10*^-/-^ HCs of both types, V_RecP_ rise times were similar (Table 2), consistent with their similar delayed rectifier profiles once K_V_1.8 was deleted (Martin et al., 2024). V_RecP_-*X* curves overlapped neatly for control and *Kcna10*^-/-^ type II HCs (**Fig. 1L**).

Aside from these expected changes, one group difference fell outside the main narrative. Resting P_O_ varied with HC type (II>I) and zone (MES>LES) independent of genotype (**Table S7**). This variation in P_O_ may reflect methodological bias (i.e. different bundle morphologies affect how the experimenter places the stiff probe on the hair bundle, see Fig 1 for examples), and should be validated with fluid-jet stimulation (Vollrath and Eatock, 2003; Peng et al., 2021).

### *Kcna10*^-/-^ HC receptor potentials had lower lowpass corner frequencies

To understand how the large differences in step-evoked responses affected frequency tuning, we displaced bundles with sinusoidal stimuli from 2 Hz to 100 Hz, spanning the upper range of natural head motion frequencies in mice (up to ∼30 Hz, Carriot et al., 2017). I_MET_ was similar across HC type and genotype (**Fig. S2**). Transduction current gain increased as stimulus frequency increased from 2 to 100 Hz, while phase shifted from a ∼20° lead to 0° (in phase with bundle displacement). Notably, both relationships were consistent with high pass filtering due to adaptation.

In contrast, V_RecP_ frequency tuning depended strongly on HC type and genotype. In control type I HCs, V_RecP_ magnitude was approximately constant across frequency (**Fig. 2A**). However, in *Kcna10*^-/-^ HCs of both types, V_RecP_ showed only lowpass tuning.

**Figure 2.**
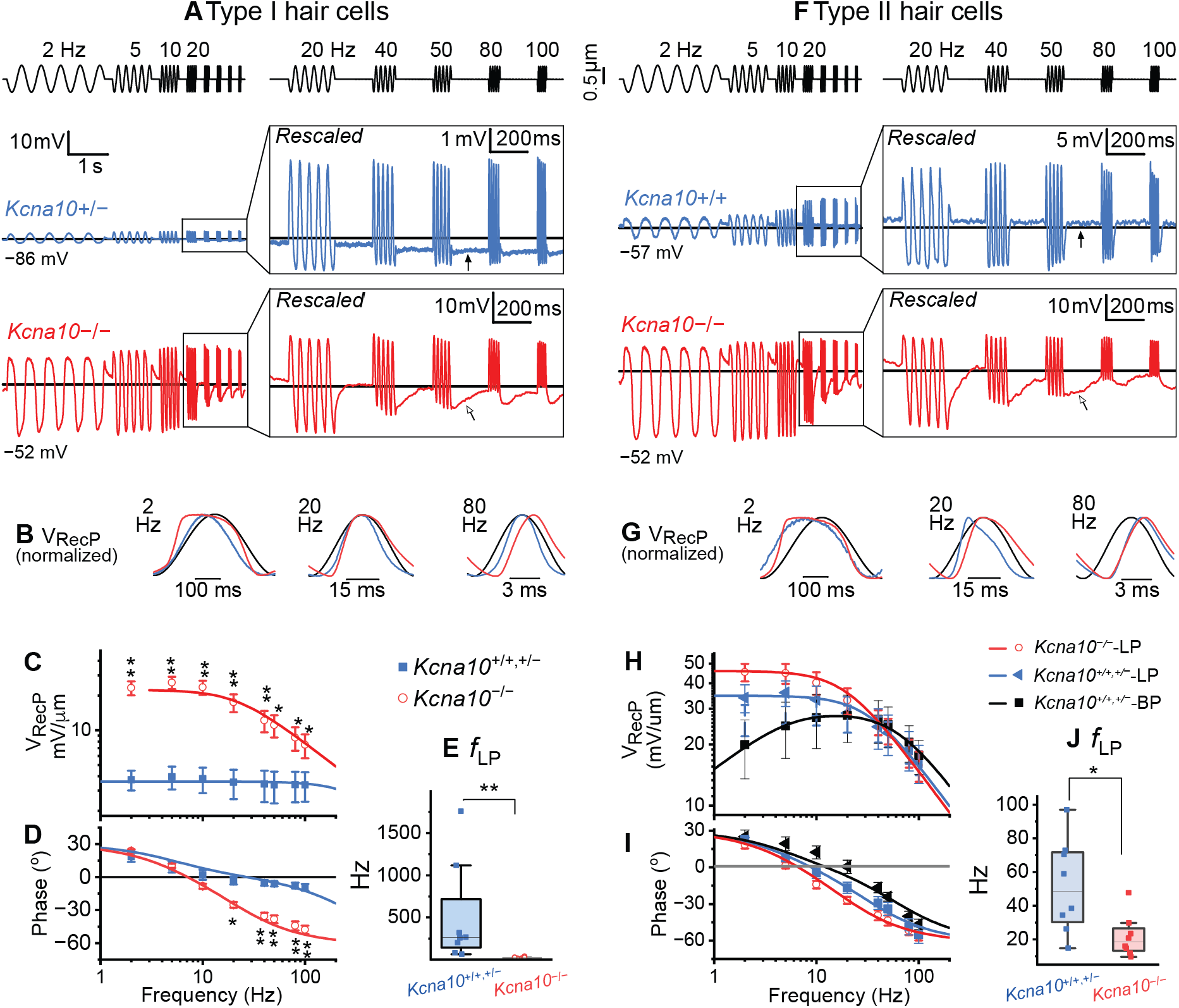
*Kcna10*^-/-^ HC receptor potentials have higher gains and lower lowpass corner frequencies than controls. (**A**) Representative type I HC responses (average of 6-13 presentations) to bundle displacement. *Left column,* bursts of sinusoid cycles from 2-100 Hz; *Right column*, bursts of sinusoid cycles from 20-100 Hz expanded in time. *Top row*, displacement; *Middle row*, *Kcna10*^+/−^ P20 LES. *Bottom row*, *Kcna10*^−/−^ P45 MES. (**B**) Normalized V_RecP_ sinusoidal cycles overlaid with the probe displacement (black). (**C-D**) Type I HC V_RecP_ referenced to X as functions of frequency; mean ± SEM values. (**C**) Peak-to-peak (AC) gain; *line*, fit to Eq. 5a. From 2-100 Hz: p = 8E-8**; 2E-8**, 1E-7**, 6E-6**, 2E-5**, 4E-4*, 0.003*, 0.008*. (**D**) Phase; *line*, fit to Eq. 5c. From 2-100 Hz: p = 0.1, 0.3, 0.2, 0.003*, 1E-6**, 3E-7**, 8E-8**, 4E-8**. (**E**) Lowpass corner frequencies (*f*_LP_) of gain-frequency relations in type I HCs. (**F**) Representative type II HC responses to bundle displacement. *Middle row*, *Kcna10*^+/+^ P22 MES. *Bottom row*, *Kcna10*^−/−^ P49 MES. (**G**) Normalized V_RecP_ sinusoidal cycle overlaid with the probe motion (black). (**H-I**) Type II HC V_RecP_ referenced to X as functions of frequency; mean ± SEM. (**H**) AC gain; fits to Eq. 5a (lowpass, LP) and Eq. 5b (bandpass, BP). Mixed ANOVA: *Kcna10*^+/+,+/−^-LP *vs Kcna10*^+/+,+/−^-BP *vs Kcna10*^−/−^-LP, _p=0.5; Frequency, p<0.0001**; Group-Frequency Interaction, p<0.0001**. # indicates a pairwise comparison between *Kcna10*^+/+,+/−^-BP and *Kcna10*^−/−^-LP. At 2Hz: p=0.02. (**I**) Phase; *line*, fit to Eq. 5c. Mixed ANOVA: Genotype, p = 0.009*; Frequency, p = 5E-26**; Genotype-Frequency Interaction, p = 0.3. At 10, 20, 40, and 50 Hz: p=0.03, 0.02, 0.01, 0.02. (**J**) Lowpass corner frequencies (*f*_LP_) of gain-frequency relations in type II HCs. *Asterisks*, uncorrected p-values from posthoc t-test: * p<0.05; ** p<0.0001.

During the 200-ms window after each sinusoid burst, control V_RecP_ stayed at a steady value close to baseline (**Fig. 2A**, black filled arrow) while *Kcna10*^-/-^ V_RecP_ had a lengthy afterhyperpolarization (**Fig 2A**, white arrow), which was likely prolonged by the slower kinetics of residual (non-K_V_1.8) g_D,R_ in *Kcna10*^-/-^ HCs (Martin et al., 2024).

As illustrated for individual cycles In **Figure 2B**, the *Kcna10*^-/-^ V_RecP_ for a single sine wave was distorted — more square than sinusoidal—at 2 Hz and lagged the stimulus >20 Hz. We quantified the response frequency dependence by fitting a sine function to each burst of 5 cycles to isolate the AC component. Despite the squarish appearance of *Kcna10*^-/-^ V_RecP_, fits to sine functions were not detectably worse than in control HCs (R^2^). The AC amplitude per unit displacement (gain) and phase relative to displacement were plotted against frequency in **Figure 2C-D**.

We estimated *f*_LP_ by fitting the gain and phase *vs.* frequency curves with a lowpass or bandpass filter transfer function (see Methods). Phase lag was much greater in *Kcna10*^-/-^ type I HCs above 10 Hz. The frequency dependence of gain and phase is summarized by the lowpass corner frequency (*f*_LP_), which fell from ∼500 Hz in control type I HCs to 22 Hz in *Kcna10*^-/-^ type I HCs (**Fig. 2E**), representing a ∼20-fold reduction in frequency bandwidth in the knockout.

Half (4/8) of control type II HCs’ V_RecP_ inherited highpass character from I_MET_ and showed overall bandpass tuning (BP), with gain increasing up until ∼40 Hz before declining (**Fig 2F-H**). The other half (4/8) showed only lowpass (LP) tuning, with gain decreasing with frequency because of filtering by the membrane time constant. In **Figure 2H-I**, *Kcna10*^+/+,+/–^-LP are displayed separately from *Kcna10*^+/+,+/–^-BP, but with combined *f*_LP_ in **Figure 2J**. The *Kcna10*^-/-^ V_RecP_ showed only lowpass tuning, with lower *f*_LP_.

In type II HCs, *f*_LP_ dropped from 52 Hz in control to 22 Hz in *Kcna10*^-/-^ type II HCs (**Fig. 2J**). In **Figure 2G**, at 20 Hz, only the control V_RecP_ had a skewed response with a prominent peak that then relaxed. This skew was observed earlier in step-evoked responses (**Fig 1K**) and other studies (Holt et al., 1999); it occurs because fast-activating g_A_ channels open and rapidly repolarize the membrane *after* V_RecP_ has charged. Despite these deviations from a classical sine waveform, fits to sine functions provided fits that were not detectably different in quality (R^2^) between *Kcna10*^-/-^ and control type II HCs.

*Kcna10*^-/-^ type I and II HCs exhibited similar lowpass tuning, reflecting their similar residual (non-K_V_1.8) delayed rectifiers (Martin et al., 2024), and consistent with the similar rise times of their step responses (**Fig. 1E.1** and **K.1** and Table 2).

Together, these whole-cell recordings show that K_V_1.8 deletion alters the gain and phase of receptor potentials relative to bundle displacement in both HC types. The reduced bandwidth and increased phase lag in *Kcna10*^-/-^ hair cells are consistent with the loss of K_V_1.8-dependent basolateral conductances and the resulting increase in input resistance (Martin et al., 2024).

### Effects of *Kcna10* deletion on calyceal voltage-dependent conductances and postsynaptic responses to type I HC hair bundle stimulation

NQ transmission from type I HCs to calyces is largely K^+^ current-driven: during a HC receptor potential, K^+^ efflux from basolateral K channels, primarily g_K,L_ (but see (Highstein et al., 2014; Sadeghi et al., 2014)), elevates cleft K^+^ (Songer and Eatock, 2013), which depolarizes the afferent through a fast electrical potential (Govindaraju et al., 2023) and a slower shift in the K^+^ equilibrium potential (E_K_) as [K^+^] rises appreciably (Lim et al., 2011; Contini et al., 2017, 2020). NQ transmission can drive afferent activity independent of glutamate-based quantal transmission (Holt et al., 2007; Pastras et al., 2023; Mukhopadhyay et al., 2026).

Based on NQ transmission’s reliance on HC K^+^ current, we asked whether NQ postsynaptic responses in calyces of *Kcna10*^-/-^ mice were affected by the absence of g_K,L_ in type I HCs. Before recording NQ, we characterized calyceal I_H_ and low-voltage-activated K^+^ (I_KLV_) current because these are proposed to contribute to NQ transmission (Contini et al., 2017, 2020). We characterized I_H_ by stepping the calyx from –64 to –124 mV and fitting an exponential to the slowly activating inward current that appears after the decay of current through deactivating K_LV_ channels (**Fig 3A**). I_H_ activated 30% more rapidly in *Kcna10*^-/-^ calyces than in control calyces (median ***τ***_Act_: 82 *vs.* 117 ms, **Fig. 3B**). To characterize I_KLV_, we measured conductance-voltage curves from tail currents (as described (Martin et al., 2024) and did not detect differences in maximal conductance (**Fig 3C**) or V_half_ (**Table 3**). We quantified calyceal excitability by recording voltage responses evoked by injected currents steps (**Fig 3D**). No differences were detected between control and *Kcna10*^-/-^ calyces in V_Rest_ (**Fig. 3E**), R_In_ (**Fig. 3F**), and spike number during depolarizing current steps (**Fig. 3G**). This suggests that for extrastriolar calyces and our recording conditions, *Kcna10* deletion had no dramatic impact on calyceal (autonomous) excitability, despite the 30% acceleration in HCN current activation.

**Figure 3.**
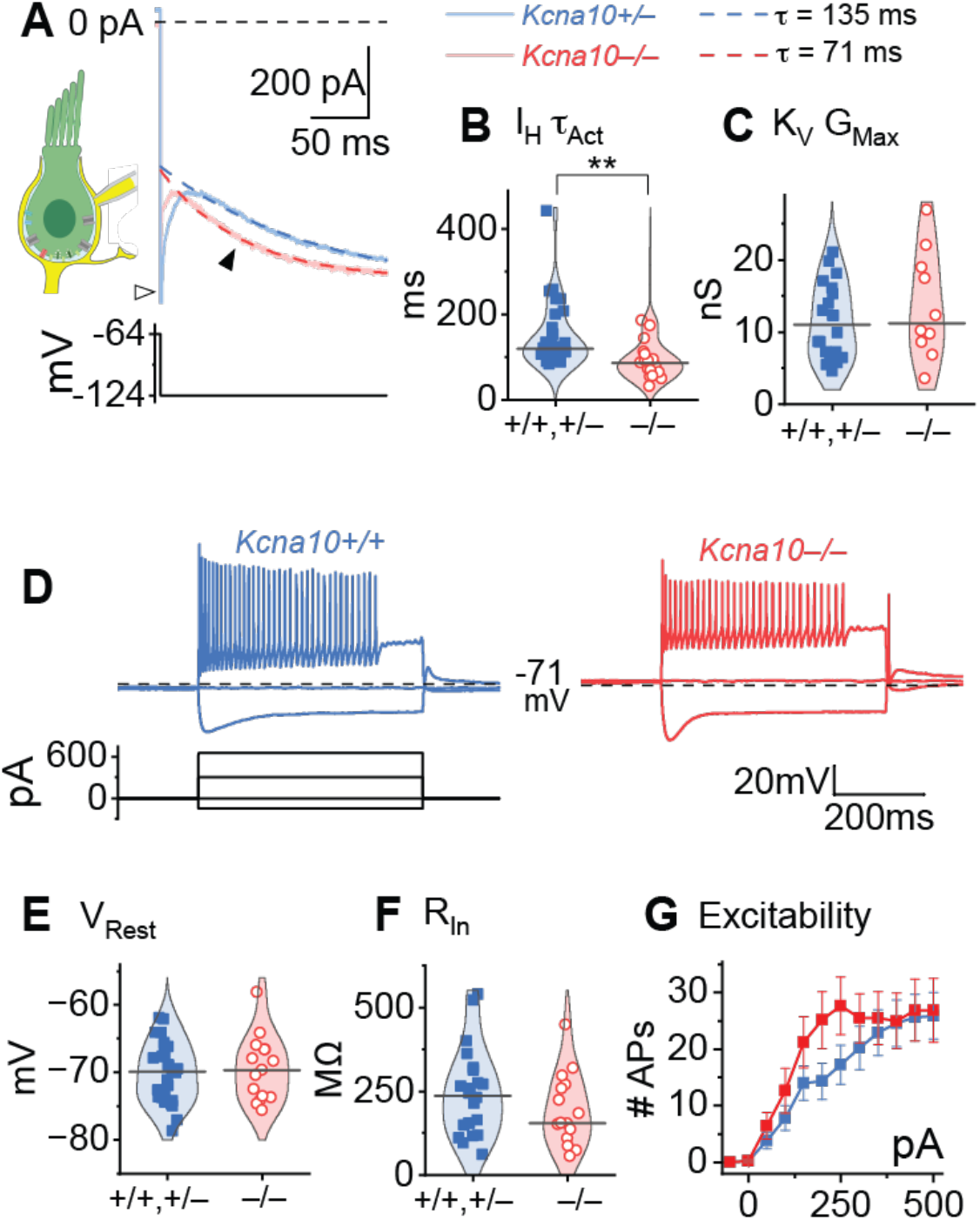
Whole-cell recordings from extrastriolar calyces showed an effect of *Kcna10* deletion on HCN current, but no detected effects on excitability. (**A**) Representative voltage-clamp recordings of I_H_ activation fit with single exponential (dashed). *Bottom panel*, holding potential. *Open arrowhead*, I_KLV_ deactivation. *Filled arrowhead*, I_H_ activation. (**B**) Time constants of exponential fits to I_H_. p=0.001 KWA, *g* 0.8. (**C**) Maximal tail conductance of K_V_ channels, showing no significant difference. (**D**) Representative current-clamp records from extrastriolar control and *Kcna10*^−/−^ calyces. *Bottom panel*, injected current. *Dashed line*, resting potential. (**E-G**) Excitability measures showed no significant differences: resting membrane potential, input resistance, and number of APs at each current injection level. Statistics in Table 3.

**Table 3.**
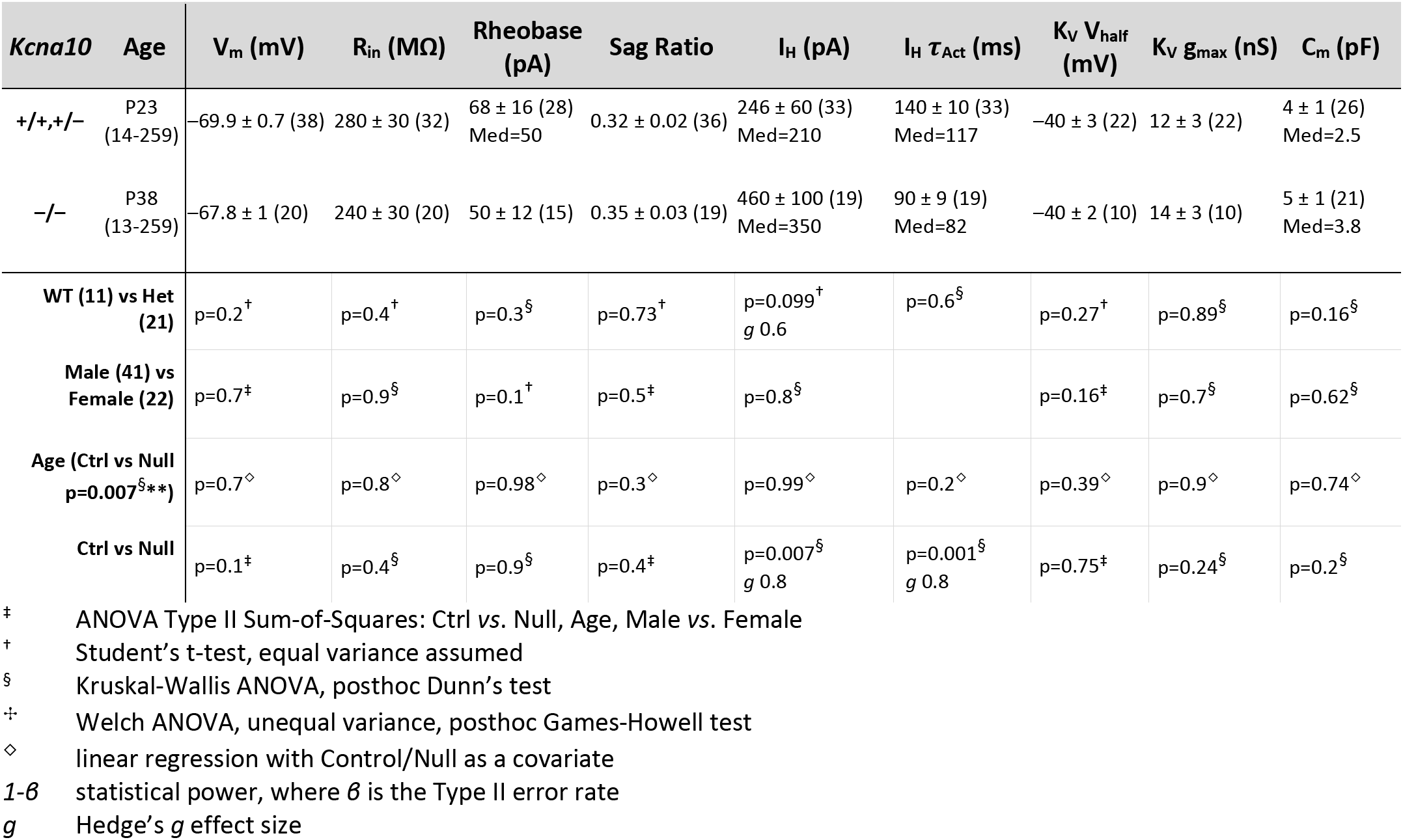
Statistics of voltage-dependent currents and excitability in utricular calyces from the extrastriola. Mean ± SEM (number of cells); median (Med) provided if data are not normally distributed (Shapiro-Wilk). Hedge’s g (*g*) provided if significant (p<0.05) or bordering on significance (p<0.1). Age is median postnatal day (range). Sag ratio is (V_SS_ – V_Min_)/(V_Rest_ – V_Min_) measured for a –200 pA injection. I_H_ amplitude and tau are parameters from exponential fits to the activation of I_H_ after hyperpolarization from –64 to –124 mV for 250 ms (R^2^>0.8).

| ***Kcna10*** | **Age** | **V<sub>m</sub> (mV)** | **R<sub>in</sub> (MΩ)** | **Rheobase (pA)** | **Sag Ratio** | **I<sub>H</sub> (pA)** | **I<sub>H</sub> τ<sub>Act</sub> (ms)** | **K<sub>V</sub> V<sub>half</sub> (mV)** | **K<sub>V</sub> g<sub>max</sub> (nS)** | **C<sub>m</sub> (pF)** |
|---|---|---|---|---|---|---|---|---|---|---|
| **+/+,+/−** | P23<br>(14-259) | −69.9 ± 0.7 (38) | 280 ± 30 (32) | 68 ± 16 (28)<br>Med=50 | 0.32 ± 0.02 (36) | 246 ± 60 (33)<br>Med=210 | 140 ± 10 (33)<br>Med=117 | −40 ± 3 (22) | 12 ± 3 (22) | 4 ± 1 (26)<br>Med=2.5 |
| **−/−** | P38<br>(13-259) | −67.8 ± 1 (20) | 240 ± 30 (20) | 50 ± 12 (15) | 0.35 ± 0.03 (19) | 460 ± 100 (19)<br>Med=350 | 90 ± 9 (19)<br>Med=82 | −40 ± 2 (10) | 14 ± 3 (10) | 5 ± 1 (21)<br>Med=3.8 |
| **WT (11) vs Het (21)** |  | p=0.2<sup>†</sup> | p=0.4<sup>†</sup> | p=0.3<sup>§</sup> | p=0.73<sup>†</sup> | p=0.099<sup>†</sup><br>*g* 0.6 | p=0.6<sup>§</sup> | p=0.27<sup>†</sup> | p=0.89<sup>§</sup> | p=0.16<sup>§</sup> |
| **Male (41) vs Female (22)** |  | p=0.7<sup>‡</sup> | p=0.9<sup>§</sup> | p=0.1<sup>†</sup> | p=0.5<sup>‡</sup> | p=0.8<sup>§</sup> |  | p=0.16<sup>‡</sup> | p=0.7<sup>§</sup> | p=0.62<sup>§</sup> |
| **Age (Ctrl vs Null p=0.007<sup>§</sup>\*\*)** |  | p=0.7<sup>◇</sup> | p=0.8<sup>◇</sup> | p=0.98<sup>◇</sup> | p=0.3<sup>◇</sup> | p=0.99<sup>◇</sup> | p=0.2<sup>◇</sup> | p=0.39<sup>◇</sup> | p=0.9<sup>◇</sup> | p=0.74<sup>◇</sup> |
| **Ctrl vs Null** |  | p=0.1<sup>‡</sup> | p=0.4<sup>§</sup> | p=0.9<sup>§</sup> | p=0.4<sup>‡</sup> | p=0.007<sup>§</sup><br>*g* 0.8 | p=0.001<sup>§</sup><br>*g* 0.8 | p=0.75<sup>‡</sup> | p=0.24<sup>§</sup> | p=0.2<sup>§</sup> |

<sup>‡</sup> ANOVA Type II Sum-of-Squares: Ctrl *vs*. Null, Age, Male *vs*. Female
<sup>†</sup> Student’s t-test, equal variance assumed
<sup>§</sup> Kruskal-Wallis ANOVA, posthoc Dunn’s test
<sup>✢</sup> Welch ANOVA, unequal variance, posthoc Games-Howell test
<sup>◇</sup> linear regression with Control/Null as a covariate
*1-β* statistical power, where *β* is the Type II error rate
*g* Hedge’s *g* effect size

We then measured NQ postsynaptic potentials by displacing the bundle of a type I HC while recording from its calyx in whole-cell patch clamp (**Fig. 4A**). Here we saw effects of *Kcna10* deletion, as expected given its strong impact on type I HC responses. Sinusoidal bundle stimulation evoked bidirectional graded postsynaptic potentials (PSPs) in the calyx (**Fig. 4B**). We typically did not observe quantal events (EPSCs or EPSPs) in calyces, consistent with other indications that quantal transmission from type I HCs becomes less and less prominent as they mature (Spaiardi et al., 2022; Michanski et al., 2023).

**Figure 4.**
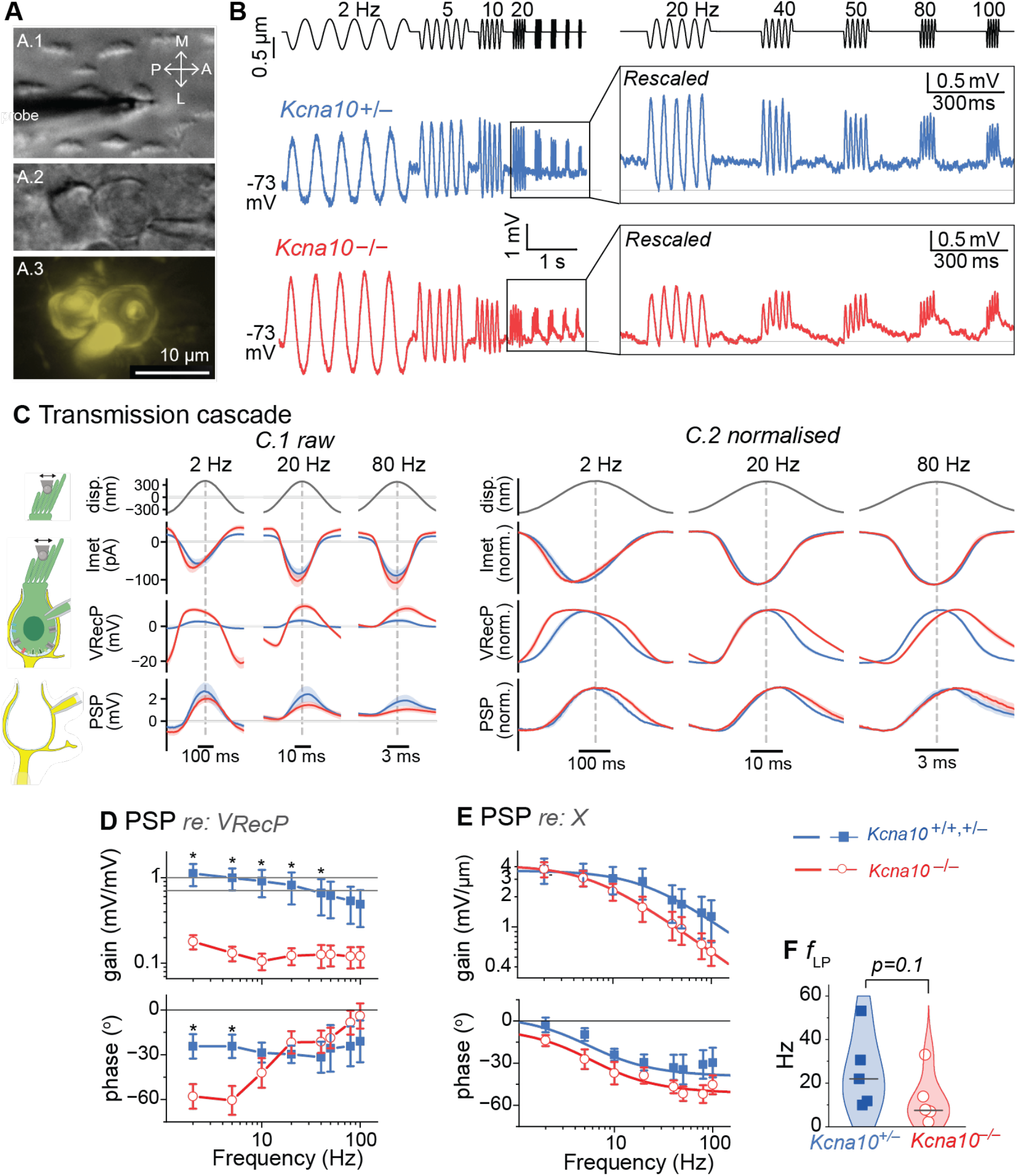
Nonquantal postsynaptic potentials show low-pass filtering with a lower corner frequency (*f*_LP_) in *Kcna10*^-/-^ calyceal afferent terminals relative to control. (**A**) Afferent arbor bearing two calyces in the lateral extrastriola; stiff probe on the bundle of the type I HC (*A.1*, DIC) enclosed in the patched calyx (*A.2*); maximum intensity projection of the dye fill (*A.3*). (**B**) Sinusoidal type I HC bundle deflections (*top, black trace*) drove voltage responses in extrastriolar calyces (averages of 4-16 presentations). (**C**) Responses to sinusoidal bundle displacement were collected from different preparations, averaged, and presented as mean ± shaded SEM for 2, 20, and 80 Hz. *C.1*, all values are referenced to baseline potential or current. *C.2*, normalized. Control HCs, n=8; *Kcna10*^-/-^ HCs, n=6; control calyces, n=5; *Kcna10*^-/-^ calyces, n=6. (**D**) Isolation of the nonquantal synaptic transfer function by referencing the population averages of calyceal and type I HC voltage responses to sinusoidal bundle displacement. Synaptic gain = V_Calyx_/V_HC_. One-tailed t-test (df=25) at 2-100 Hz: p=0.008*, 0.003*, 0.02*, 0.02*, 0.04*, 0.05, 0.06, 0.06. Synaptic phase = φ_Calyx_ – φ_HC_. One-tailed t-test (df=25) at 2-100 Hz pairwise 2: p=0.004*, 0.004*, 0.1, 0.8, 0.8, 0.7, 0.8, 0.8. (**E**) Calyceal postsynaptic potential gain and phase referenced to the bundle displacement, fit to Eq. 5a (gain) and 5d (phase). Mixed ANOVA for gain: Genotype, p=0.5; Frequency, p=1E-18; Genotype-Frequency Interaction, p=0.4. NQ phase. Mixed ANOVA for phase: Genotype, p=0.1; Frequency, p=0.001; Genotype-Frequency Interaction, p=0.6. (**F**) Lowpass corner frequencies of PSP gain-frequency curves in *E*. *Asterisks*: *, p<0.05; **, p<0.01. Posthoc p-values and fit parameters can be found in Table 4.

**Figure 4C** illustrates how the input sinusoid is altered during each stage of signal transmission: from transduction currents to receptor potentials to postsynaptic potentials. All traces, which are population averages of type I HC and calyx recordings (Table 4), are shown at their original scale in C.1 and then normalized in C.2 to emphasize how the sinusoidal waveform is shaped both by K_V_1.8 deletion and by the transmission cascade. The transduction current peaked in advance of bundle displacement for both control and *Kcna10*^-/-^ HCs. The receptor potential peaked in advance at 2 Hz, again for both, but was shifted more negatively at higher frequencies, with the largest effect in the *Kcna10*^-/-^ HCs. For the calyx NQ PSP, further slowing put the phase difference at about 0 at 2 Hz and negative (a phase lag) at 20 and 80 Hz, for both control and *Kcna10*^-/-^ calyces. At 2-20 Hz, *Kcna10*^-/-^ PSPs overlapped with control PSPs during the ascending (depolarizing) phase, but lagged during the descending (hyperpolarizing) phase.

**Table 4.**
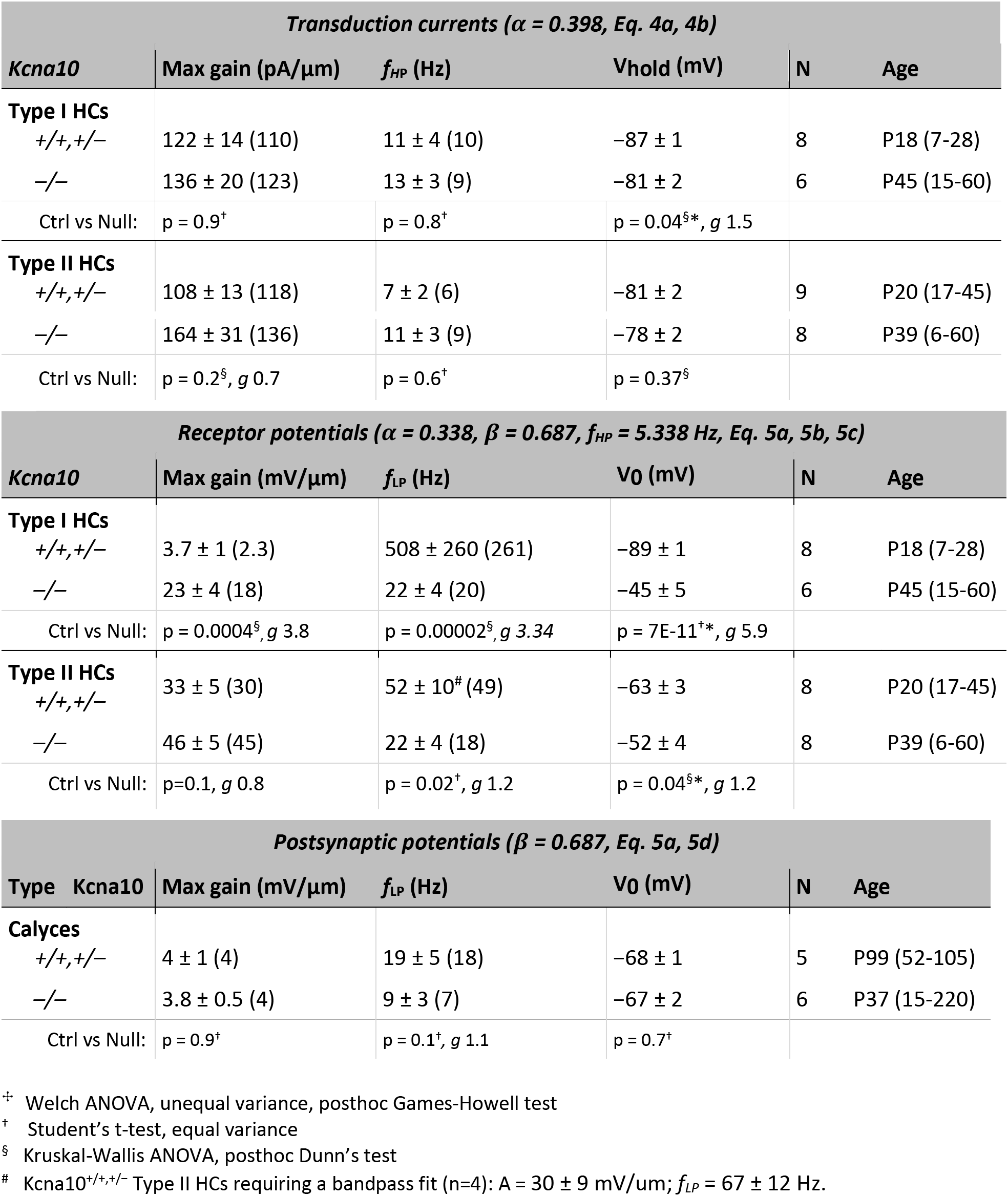
Frequency tuning of transduction currents, receptor potentials, and postsynaptic potentials. V_0_ was measured from the first 200 ms of the recording. We report mean ± SEM (range). We report the p-value, effect size (Hedge’s *g*) if p ≤ 0.1, and statistical power (1-β) if p > 0.05. Sample size (N) is number of cells. Age is median postnatal day (range).

| *Transduction currents (α = 0.398, Eq. 4a, 4b)* |  |  |  |  |  |
|---|---|---|---|---|---|
| *Kcna10* | Max gain (pA/µm) | *f<sub>HP</sub>* (Hz) | Vhold (mV) | N | Age |
| **Type I HCs** |  |  |  |  |  |
| *+/+,+/–* | 122 ± 14 (110) | 11 ± 4 (10) | −87 ± 1 | 8 | P18 (7-28) |
| *–/–* | 136 ± 20 (123) | 13 ± 3 (9) | −81 ± 2 | 6 | P45 (15-60) |
| Ctrl vs Null: | p = 0.9<sup>†</sup> | p = 0.8<sup>†</sup> | p = 0.04<sup>§</sup>\*, *g* 1.5 |  |  |
| **Type II HCs** |  |  |  |  |  |
| *+/+,+/–* | 108 ± 13 (118) | 7 ± 2 (6) | −81 ± 2 | 9 | P20 (17-45) |
| *–/–* | 164 ± 31 (136) | 11 ± 3 (9) | −78 ± 2 | 8 | P39 (6-60) |
| Ctrl vs Null: | p = 0.2<sup>§</sup>, *g* 0.7 | p = 0.6<sup>†</sup> | p = 0.37<sup>§</sup> |  |  |
| *Receptor potentials (α = 0.338, β = 0.687, f<sub>HP</sub> = 5.338 Hz, Eq. 5a, 5b, 5c)* |  |  |  |  |  |
| *Kcna10* | Max gain (mV/µm) | *f*<sub>LP</sub> (Hz) | V0 (mV) | N | Age |
| **Type I HCs** |  |  |  |  |  |
| *+/+,+/–* | 3.7 ± 1 (2.3) | 508 ± 260 (261) | −89 ± 1 | 8 | P18 (7-28) |
| *–/–* | 23 ± 4 (18) | 22 ± 4 (20) | −45 ± 5 | 6 | P45 (15-60) |
| Ctrl vs Null: | p = 0.0004<sup>§</sup>, *g* 3.8 | p = 0.00002<sup>§</sup>, *g 3.34* | p = 7E-11<sup>†</sup>\*, *g* 5.9 |  |  |
| **Type II HCs**<br>*+/+,+/–* | 33 ± 5 (30) | 52 ± 10<sup>#</sup> (49) | −63 ± 3 | 8 | P20 (17-45) |
| *–/–* | 46 ± 5 (45) | 22 ± 4 (18) | −52 ± 4 | 8 | P39 (6-60) |
| Ctrl vs Null: | p=0.1, *g* 0.8 | p = 0.02<sup>†</sup>, *g* 1.2 | p = 0.04<sup>§</sup>\*, *g* 1.2 |  |  |
| *Postsynaptic potentials (β = 0.687, Eq. 5a, 5d)* |  |  |  |  |  |
| Type Kcna10 | Max gain (mV/µm) | *f*<sub>LP</sub> (Hz) | V0 (mV) | N | Age |
| **Calyces** |  |  |  |  |  |
| *+/+,+/–* | 4 ± 1 (4) | 19 ± 5 (18) | −68 ± 1 | 5 | P99 (52-105) |
| *–/–* | 3.8 ± 0.5 (4) | 9 ± 3 (7) | −67 ± 2 | 6 | P37 (15-220) |
| Ctrl vs Null: | p = 0.9<sup>†</sup> | p = 0.1<sup>†</sup>, *g* 1.1 | p = 0.7<sup>†</sup> |  |  |

<sup>✢</sup> Welch ANOVA, unequal variance, posthoc Games-Howell test
<sup>†</sup> Student’s t-test, equal variance
<sup>§</sup> Kruskal-Wallis ANOVA, posthoc Dunn’s test
<sup>#</sup> Kcna10<sup>+/+,+/–</sup> Type II HCs requiring a bandpass fit (n=4): A = 30 ± 9 mV/um; *f<sub>LP</sub>* = 67 ± 12 Hz.

As with HCs, we fit a sine function to each burst of 5 cycles to isolate the AC component (goodness-of-fit: R^2^ > 0.5). To compare presynaptic and postsynaptic signals directly, we referenced the population average of calyx NQ PSPs to the population average of type I HC V_RecP_s. These values describe the synaptic transfer function. Synaptic gain was near unity in controls, indicating efficient, 1:1 transfer of the presynaptic voltage to the calyx (**Fig. 4D**). In contrast, *Kcna10*^-/-^ synapses had much lower gain such that much larger V_RecP_ were required to achieve the same postsynaptic depolarization. (Recall that *Kcna10*^-/-^ type I HCs have a larger and slower V_recP_ because R_in_ increases substantially when K_V_1.8 is deleted (**Fig. 2A,C**).) For control data, phase of the calyx NQ PSP relative to the HC V_RecP_ stayed ∼–30° across frequencies (**Fig 4D**). In *Kcna10*^-/-^ calyces, phase was much more frequency-dependent, ranging from ∼–60° below 10 Hz to control-like values at higher frequencies (∼–30°), an effect evident in the time domain at 2 and 20 Hz in **Figure 4C**.

Referencing PSPs to the displacement stimulus, the low-frequency (2 Hz) AC gain was similar for control and *Kcna10*^-/-^ calyces (∼4 mV/μm, **Fig 4E**), but median *f*_LP_ dropped from 19 Hz (range 6-34 Hz, n=5) in control calyces to 9 Hz (1-24 Hz, n=6) in *Kcna10*^-/-^ calyces (p=0.1, Kruskal-Wallis ANOVA with effect size (Hedge’s g) of 1.1). This change in NQ transmission is expected from presynaptic slowing and qualitatively consistent with the fast component of NQ transmission as previously modeled (Govindaraju et al., 2023).

Note that PSP gains referenced to displacement are likely larger *in vivo* where they usually contribute to a common afferent arbor that simultaneously integrates inputs from multiple type I and II HCs. However, in our *ex vivo* experiments, single bundle displacements rarely evoked action potentials; in the extrastriola, NQ PSPs are usually diluted across a relatively large extrastriolar arbor with a distant spike initiation zone that integrate NQ and quantal inputs from multiple HCs. In a similar semi-intact preparation of the rat saccule, with similar stimulation methods, NQ PSPs in striolar calyces did evoke spikes (Songer and Eatock, 2013; their Fig. 5). Striolar afferents have long been described as more sensitive to inputs, both sensory and efferent (Goldberg, 2000); possible factors are that the striolar type I HCs have more g_K,L_ (K_V_1.8) channels to contribute to NQ transmission, and striolar calyces belong to more compact terminal arbors with a shorter distance from the calyx postsynaptic membrane to the spike initiation zone (Lysakowski et al., 2011).

**Figure 5.**
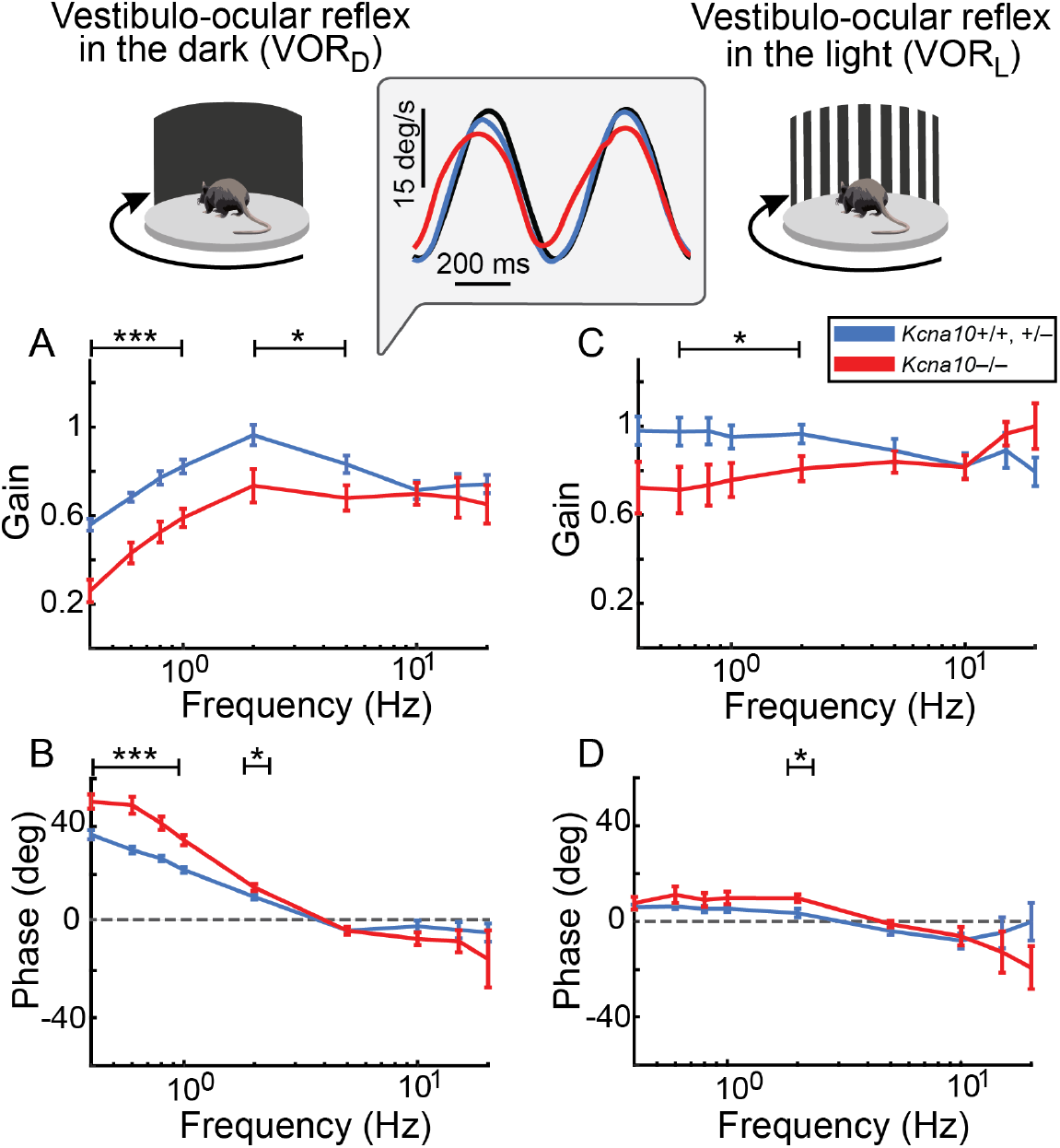
Horizontal VOR differs between *Kcna10*^−/−^ and control mice. (**A, B**) Vestibulo-ocular reflex in the dark (VOR_D_) data for *Kcna10*^+/+,+/−^ mice (blue, n = 14, aged 10-20 months, median 18) are compared with *Kcna10*^−/−^ mice (red, n = 9, aged 10-20 months, median 19). (**C, D**) Vestibulo-ocular reflex in the light (VOR_L_) data for *Kcna10*^+/+,+/−^ mice (blue, n = 12, aged 10-20 months, median 18) are compared with *Kcna10*^−/−^ mice (red, n = 9, aged 10-20 months, median 19). **Inset:** Example traces of head velocity (black) and eye velocity for *Kcna10*^+/+,+/−^ mice (blue) and *Kcna10*^−/−^ mice (red) at 1 Hz in the dark. Eye velocities were inverted for comparison with head velocities. Mean ± SEM. *Asterisks*: *p < 0.05, ***p < 0.001.

## II. Vestibulomotor behaviors

We next assessed whether the lack of K_V_1.8 affected performance during vestibular-associated tasks in mice aged 2-12 months (**Tables S2-S4**). *Kcna10*^+/+^ and *Kcna10*^+/–^ mice were pooled as controls when there was no detectable difference between them on a given task. *Kcna10*^-/-^ mice were healthy, developed normally, and did not display overt signs of vestibular dysfunction such as circling and head tilt, as noted previously (Lee et al., 2013). K_V_1.8 is not expressed in neural tissue, but, given that Lee et al. (2013) reported expression in skeletal muscle (and kidney), we first characterized behavior on general motor and motor learning tasks. Both *Kcna10*^-/-^ and control mice successfully learned a rotarod task in the light. After training, latencies to fall were similar across genotype: 80 ± 7 s (*Kcna10^+/–^*; 3 males, 7 females; age 2-6.9 months, median 3.8) and 82 ± 9 s (*Kcna10^-/-^*; 5 males, 5 females; age 2-6.9 months, median 3.8; ANOVA Tukey’s p=0.85, power 0.05). Control and *Kcna10*^-/-^ mice did not walk in a detectably different manner, which we assessed by quantifying gait properties, such as stance width and paw angle, during a forced treadmill run (**Table 5**). Grip strength was also similar across genotype: *Kcna10^+/–^*, 184 ± 9 g, n=19; and *Kcna10^-/-^*, 173 ± 8 g, n=20; 2-way ANOVA: Genotype, Tukey’s p=0.36, power 0.16; Sex, Tukey’s p=0.003, g 1.36).

**Table 5.** Gait parameters of control (*Kcna10^+/+,+/–^*) and *Kcna10^-/-^* mice during forced treadmill running at fast and slow speeds (25 and 15 cm/s). Values are mean ± SEM across mice. Sample sizes are n=16 control mice (aged 10-30 weeks, median 12) and n=12 *Kcna10^-/-^* mice (aged 10-30 weeks, median 12). Statistics are from a linear mixed model (genotype x limb x speed, with sex and age as covariates and mouse as a random intercept); p-values are main effects, averaged over the other factors. Age is reported as the model slope per week. Effect sizes are Hedges’ g, unadjusted for covariates.

|  | **Paw Angle (deg)** | **Stance Width (mm)** | **Stride Dur (ms)** | **Swing Dur (ms)** | **Stance Dur (%)** | **Stride Length (mm)** | **Stride Length Variability** |
|---|---|---|---|---|---|---|---|
| **Fast** |  |  |  |  |  |  |  |
| *Ctrl* |  |  |  |  |  |  |  |
| Fore | 22.3 ± 0.6 | 7.45 ± 0.2 | 257 ± 5.2 | 97.4 ± 2.6 | 62.2 ± 0.97 | 64.2 ± 1.3 | 1.01 ± 0.06 |
| Hind | 25.4 ± 1.1 | 13.2 ± 0.3 | 260 ± 4.4 | 61.7 ± 4.5 | 77 ± 1.7 | 65 ± 1.1 | 1.25 ± 0.13 |
| *Null* |  |  |  |  |  |  |  |
| Fore | 21.7 ± 0.6 | 7.65 ± 0.15 | 254 ± 4.4 | 89.2 ± 3.1 | 65.2 ± 1.3 | 63.4 ± 1.1 | 1.13 ± 0.08 |
| Hind | 24 ± 0.99 | 12.9 ± 0.27 | 258 ± 4.7 | 56.8 ± 3.2 | 79.8 ± 1 | 64.4 ± 1.2 | 1.54 ± 0.16 |
| **Slow** |  |  |  |  |  |  |  |
| *Ctrl* |  |  |  |  |  |  |  |
| Fore | 20.4 ± 0.53 | 7.7 ± 0.2 | 349 ± 7.5 | 111 ± 3.8 | 68.4 ± 0.8 | 52.4 ± 1.1 | 1.1 ± 0.1 |
| Hind | 28.5 ± 1.2 | 13.3 ± 0.3 | 354 ± 8.3 | 62.9 ± 5.2 | 86.7 ± 2.1 | 53.1 ± 1.2 | 1.5 ± 0.1 |
| *Null* |  |  |  |  |  |  |  |
| Fore | 21 ± 0.6 | 8.1 ± 0.1 | 351 ± 8.2 | 106 ± 3.9 | 70.9 ± 1 | 52.6 ± 1.2 | 1.2 ± 0.1 |
| Hind | 27.4 ± 1.1 | 13 ± 0.3 | 354 ± 8.5 | 64 ± 3.6 | 85.5 ± 1.6 | 53.1 ± 1.3 | 1.5 ± 0.1 |
| Fore vs Hind | <0.001;<br>g=1.6 | <0.001;<br>g=+6.4 | 0.2;<br>g=+0.2 | <0.001;<br>g=-2.8 | <0.001;<br>g=+3 | 0.1;<br>g=+0.2 | <0.001;<br>g=+1 |
| Fast vs Slow | 0.08;<br>g=+0.4 | 0.08;<br>g=+0.3 | <0.001;<br>g=+3.8 | <0.001;<br>g=+0.7 | <0.001;<br>g=+1.6 | <0.001;<br>g=-2.5 | 0.1;<br>g=+0.4 |
| Age (wk<sup>-1</sup>) | 0.02;<br>+0.175/wk | 0.5;<br>+0.018/wk | 0.12;<br>-1.21/wk | 0.010;<br>+1.02/wk | <0.001;<br>-0.5/wk | 0.2;<br>-0.21/wk | 0.9;<br>-0.0013/wk |
| M vs F | 0.7;<br>g=-0.1 | 0.7;<br>g=+0.07 | 0.13;<br>g=+0.6 | 0.718;<br>g=-0.14 | 0.9;<br>g=+0.3 | 0.14;<br>g=+0.7 | 0.7;<br>g=+0.2 |
| Ctrl vs Null | 0.3;<br>g=-0.2 | 0.84;<br>g=-0.02 | 0.7;<br>g=+0.1 | 0.2;<br>g=-0.3 | 0.08;<br>g=+0.5 | 0.7;<br>g=-0.2 | 0.3;<br>g=+0.5 |

### Vestibulo-ocular reflexes in *Kcna10*^-/-^ mice have reduced gain and more variable phase

HC-afferent synapses are the first in the VOR relay from HC to afferent to vestibular nucleus to motor nuclei to eye muscles. The VOR stabilizes gaze by moving the eye in opposition to the ongoing head motion. In rhesus monkeys, the horizontal angular VOR can have a latency as short as 5-6 ms (Huterer and Cullen, 2002). Given the effects of K_V_1.8 on HC response timing (phase) and gain (which independently affects the speed at which afferent spikes are triggered), we looked for effects on VOR. We measured horizontal angular VOR responses in head-fixed *Kcna10*^−/−^ mice and their control counterparts by rotating the mouse horizontally while tracking pupil movement with a platform-mounted camera. VORs were measured in the light (VOR_L_), where vision contributes, and the dark (VOR_D_), where visual input is absent, thereby isolating vestibular-driven responses.

VOR_D_ came closer in control mice than *Kcna10*^−/−^ mice to ideal values for gain (1) and phase (0°) over much of the frequency range tested: <10 Hz for gain and <5 Hz for phase (per-frequency statistics in Table S1). The Fig. 5 inset shows example head- and eye-velocity traces at 1 Hz in the dark, illustrating the control-*Kcna10*^−/−^ difference. *Kcna10*^−/−^ mice also had lower VOR_L_ gain (**Fig. 5C**), as noted in another model of vestibular dysfunction (Giese et al., 2025).

### *Kcna10*^-/-^ mice have specific deficits on challenging balance tasks

In an open arena, *Kcna10*^-/-^ mice exhibited normal centrophobism and overall activity but were less likely to rear up on their hindlegs (**Fig. 6A, Table S2**). Reduction in this normal exploratory behavior likely reflects impaired postural stability, not heightened anxiety, which would have been revealed as higher centrophobism levels.

**Figure 6.**
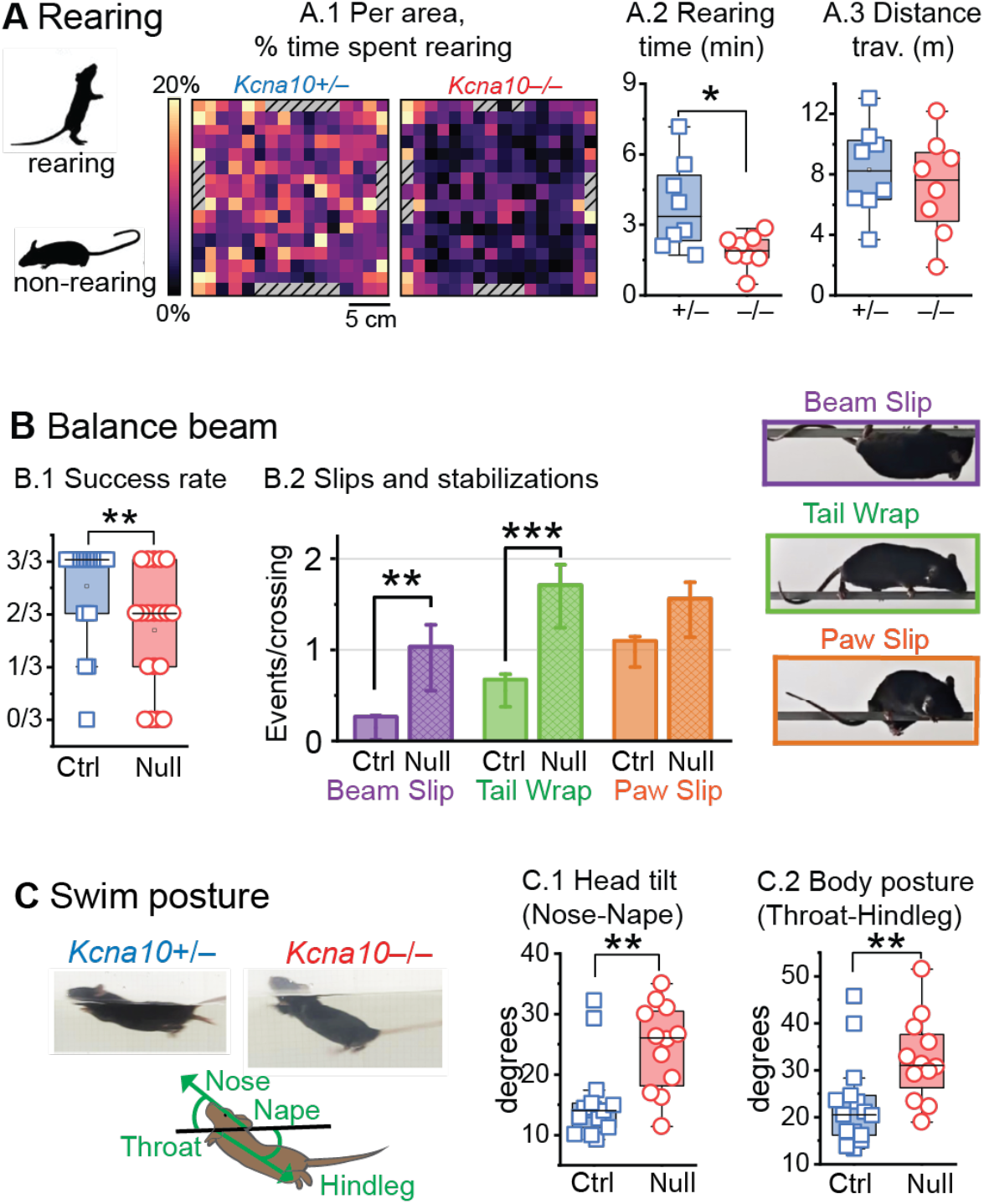
*Kcna10*^-/-^ mice show instability in challenging balance tasks and have altered swim posture. (**A**) 1-hr exploration of an arena. (A.1) Heatmap of percent of time spent rearing in each 2.56 cm^2^ floor square, averaged across by *Kcna10*^+/−^ (N=8) and *Kcna10*^−/−^ (N=8) mice. Grey, patterned squares lack data. (A.2) Time spent rearing (p=0.02 Welch ANOVA, g 1.4). (A.3) Total distance traveled (p=0.5 ANOVA). (**B**) Crossing a narrow (6-mm wide) balance beam by *Kcna10*^+/+,+/−^ mice (N=24) and *Kcna10*^−/−^ mice (N=19). (B.1) Fraction of trials on which mice crossed without requiring rescue (p=0.002 KWA, g 0.9) (B.2) Per trial occurrences of beam slips, tail wraps, and paw slips per mouse. (**C**) Body posture and head tilt during a 30-second swim in *Kcna10*^+/+,+/−^ (n=15) and *Kcna10*^-/-^ (n=9) mice. (C.1) Head pitch measured as the angle between the nose-nape line and the waterline. (C.2) Body posture measured as the angle between the throat-hindleg line and the waterline. Asterisks: * p<0.05, ** p<0.01.

Most control mice (86%) but only half of *Kcna10*^-/-^ mice (56%) mice crossed a narrow (6 mm) balance beam without slipping off the beam (p<0.01, **Fig. 6B**). *Kcna10*^-/-^ mice frequently wrapped their tails around the beam for additional stability (**Fig 6B.2**) and were slower to cross, but not significantly so (14 ± 2 *vs.* 20 ± 4 s, p<0.1, *g* 0.5, **Suppl. Table 3**).

Forced swimming tests can help identify vestibulo-motor defects by reducing stabilizing proprioceptive contact with the ground (Sawada et al., 1994; Jones et al., 2004). We found that *Kcna10*^-/-^ mice stayed afloat but swam less effectively than controls, with a greater angle (tilt) relative to the water surface (**Fig. 6C**, see Supplemental Video). We quantified head tilt and body tilt by measuring the angles between water line and either the nape-nose line (for head tilt) or the throat-hindleg line (for body tilt; see diagram in **Fig. 6C**). A nape-nose angle near 0° means the head lies perfectly parallel to the water surface, and a throat-hindleg angle near 0° means the body is floating parallel to the water surface. Both nape-nose and throat-hindleg angles were smaller by about ∼10° in controls compared to *Kcna10*^-/-^ mice.

When possible, we tested vestibulomotor outcomes for sex differences. In the open field arena, male mice spent more time rearing than females (3.6 ± 0.7 *vs.* 2.1 ± 0.4 min, p=0.02, *g* 1.0, **Suppl. Table 2**).

### *Kcna10*^-/-^ mice have altered head motions across multiple balance tasks

We quantified head control with head-mounted inertial measurement units (IMU) and compared the resulting power spectra between control and *Kcna10*^-/-^ animals. During spontaneous exploration of the open field arena, *Kcna10*^-/-^ mice had lower head motion power in the fore-aft, vertical, and pitch dimensions (**Fig. 7A**). This reduction accompanied less rearing, although overall locomotor activity was similar between groups (**Fig. 6A, Table S2**). The lower head motion power observed in the open field was not evident on the balance beam. Instead, on the beam, *Kcna10*^-/-^ mice had slightly higher head motion power at the higher frequencies across all dimensions (**Fig. 7B**).

**Figure 7.**
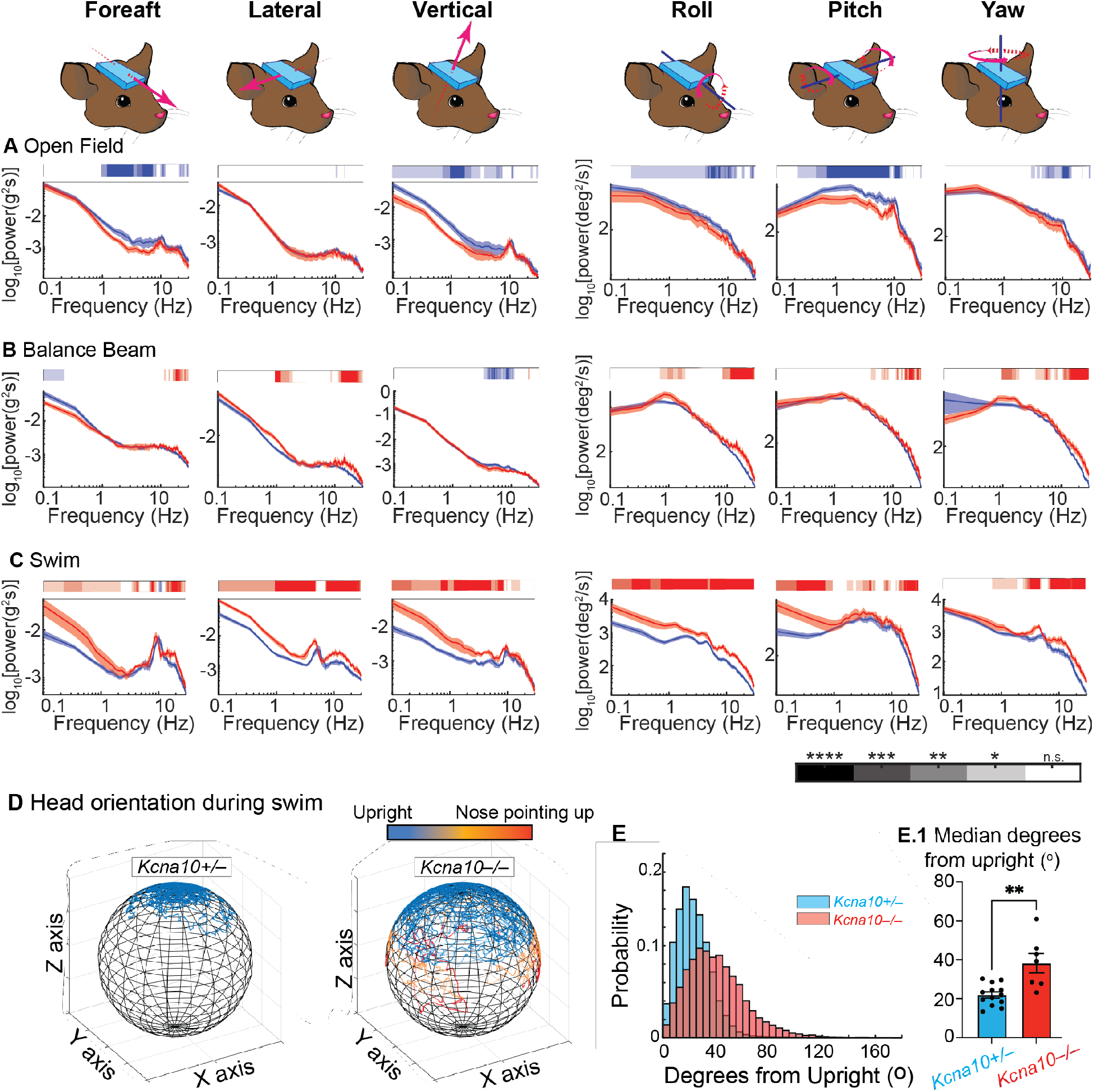
Instrumented head motion detection shows altered head motions and orientation in *Kcna10*^-/-^ mice. Cartoons show head-mounted IMU device detecting head motions along the three translational and three rotational axes: from left to right, Foreaft, Lateral, Vertical, and Roll, Pitch, Yaw. (**A-C**) Head motion power spectra were recorded along each of the 6 axes during three behavioral settings. Mean ± shaded SEM. Blue bars above data show frequency ranges where *Kcna10*^+/+,+/−^ mice had significantly greater head motion power, and red bars mark ranges where *Kcna10*^−/−^ mice had significantly greater head motion power. The intensity of blue and red bars corresponds to levels of significance on the color map at the bottom of C. Data below the noise floor is not shown. (**A**) Passive exploration of an open field arena in *Kcna10*^+/+,+/−^ (n=11) and *Kcna10*^−/−^ (n=7) mice. (**B**) Crossing a narrow balance beam in *Kcna10*^+/+,+/−^ (n=11) and *Kcna10*^−/−^ (n=6) mice. (**C**) 1-minute forced swim in *Kcna10*^+/+,+/−^ (n=13) and *Kcna10*^−/−^ (n=7) mice. (**D**) Spherical representations of head orientation relative to gravity for a representative control (left) and *Kcna10*^-/-^ (right) mouse. IMU head vector pitched to be aligned with gravity (upright) as described in Hughes et al. 2024. Data colorized to reflect deviation from upright: blue = deviations 0°-65°, orange = deviations 65°-90°, red = deviations >90°. (**E**) Histogram of individual swim trials (n=3 per mouse) for control (blue, N=13) and *Kcna10*^-/-^ (red, N=7) mice. Box plot shows the median (black dot), interquartile range (filled rectangle), and whiskers corresponding to approximately ± 2.7 standard deviations. (E.1) Median degrees from upright per mouse, p=0.0047, Mann-Whitney test. Mean ± SEM shown. *Asterisks*: * p<0.05, ** p<0.01, *** p<0.001, ****p<0.0001.

Swimming revealed the largest differences in head motion power spectra between control and *Kcna10*^−/−^ mice. While swimming, *Kcna10*^-/-^ mice had greater head motion power in all dimensions and over a wide range of frequencies (**Fig. 7C**). These increases may reflect impaired head stabilization resulting from slower vestibular signaling in *Kcna10*^-/-^ mice. In addition, IMU-based estimates of head orientation corroborated the differences observed in the video-based measurements during swimming. The forehead vector was closer to the upright vertical axis in controls than *Kcna10*^-/-^ mice (approximately 20 *versus* 40° degrees from upright, **Fig 7D**).

Overall, vestibulomotor testing suggests that K_V_1.8 contributes to gaze and postural stability, particularly during challenging tasks such as swimming and crossing narrow balance beams. Together with the electrophysiological findings, these results support a role for basolateral HC K^+^ conductances in shaping afferent signals that support vestibulomotor control.

## Discussion

Our comparisons show that for both HC types in the extrastriolar mouse utricle, K_V_1.8 attenuates and speeds up receptor potentials, as predicted from effects of K_V_1.8 on potentials evoked by injected currents (Martin et al., 2024). Further, by comparing control type I HC receptor potentials with control calyx NQ PSPs for similar bundle stimuli, we show that the low gain of the type I HC receptor potential is compensated by the high synaptic transfer gain (PSP re: receptor potential) of NQ transmission. The effects of K_V_1.8 subunits in HCs may largely account for the impairment of vestibulomotor behaviors that we documented in constitutive knockouts, especially given that K_V_1.8 subunits are strongly expressed in HCs and not in neurons.

### K_V_1.8 and signal processing (receptor potentials and synaptic transmission)

Control type I HC receptor potentials had low gains and fast membrane charging times, consistent with the low input resistance imparted by large numbers of g_K,L_ channels with their hyperpolarized activation range (Correia, 1992; Chen and Eatock, 2000; Hurley et al., 2006; Dhawan et al., 2010; Contini et al., 2012). Small receptor potentials have little effect on the activation of voltage-dependent channels, making the receptor potential more linear and less dynamic.

These findings extend our previous current injection experiments (Martin et al., 2024) to the more naturalistic setting of displacement-evoked receptor potentials, where the input is the nonlinear MET current. The present approach reveals that V_RecP_ saturates at smaller displacements in *Kcna10*^-/-^ type I HCs than in control type I HCs. This is readily explained by the larger displacement-evoked depolarizations activating more delayed rectifier K_V_ channels, which oppose further depolarization. In contrast, g_K,L_ attenuates the voltage changes, centering the linear region of the sigmoidal V_RecP_-X curve at larger (∼0.4 micron) displacements, without affecting slope factor.

In the frequency domain, fast membrane charging times gave control type I HC receptor potentials a flat, frequency-independent gain up to 100 Hz (**Fig. 2C)**, with phase close to 0° throughout (**Fig. 2D**). The difference in these relationships between control and *Kcna10*^-/-^ HC data (**Fig. 2**) was similar to the difference for maturing type I HCs with and without g_K,L_ in the striolar zone of the rat saccule, tested as g_K,L_ (now known to be a K_V_1.8 conductance) was acquired over the first postnatal week (Songer and Eatock, 2013).

Owing to adaptation of I_MET_ and fast activation of g_A_, control type II HCs show a rapid receptor potential decay during step displacements and at 5-20 Hz, as noted by others (Holt et al., 1999), and faster membrane charging times (*τ*_Onset_) from V_Rest_. *τ*_Onset_ measured with bundle deflection were 3-5-fold shorter than those measured with current injection (Martin et al., 2024), in *Kcna10^-/-^*type I HCs, control type II HCs, and *Kcna10^-/-^* type II HCs (all p < 0.02). During a true V_RecP_, membrane charging time is actively reduced as MET channels open and lower R_In_.

Biophysical modeling predicts that g_K,L_ contributes to NQ transmission at the type I HC-calyx synapse by modulating fast electrical and slow K^+^ potentials (Govindaraju et al., 2023). We show that *Kcna10*^-/-^ HCs also support NQ transmission, although with slower dynamics. Modeling suggests this difference is attributable to *Kcna10*^-/-^ type I HCs’ higher R_in_ plus the time required for the residual non-*K_V_1.8* channels to open during a depolarizing receptor potential. In controls, g_K,L_ (K_V_1.8) confers low membrane resistance, which allows transduction current to flow directly into resistive current (through g_K,L_ channels into the cleft, to NQ) instead of entirely into capacitive current (charging the receptor potential).

As a result, the presence of K_V_1.8 reduces (1) phase lag and (2) distortion (nonlinearity) of peak HC RP and calyx PSP referenced to hair bundle displacement, as shown with normalized data in **Fig. 4C.2**. With non-normalized data, K_V_1.8 is seen to strongly reduce the gain of type I HC receptor potential re: bundle displacement (**Fig. 2C, 4C.1**) without affecting the gains of PSP re: displacement. This is achieved by a remarkable increase in the gain of the calyx PSP re: type I HC receptor potential (Fig. 4D). This last observation highlights how the presence of K_V_1.8 improves response fidelity by increasing speed and linearity. Adding low-voltage activated K_V_ channels is expected to increase speed, but at a cost in signal gain (Rothman and Manis, 2003). Here, the cost is avoided by the unusual NQ transmission.

I_H_ in calyces can carry NQ currents (Contini et al., 2017), raising the possibility that the faster-activating I_H_ in *Kcna10*^-/-^ calyces is compensatory. Because I_H_ is cAMP-sensitive and potentially subject to efferent modulation (Bronson and Kalluri, 2023) and extracellular [K^+^] (Meredith et al., 2012), it is a plausible substrate for signaling homeostasis.

Our observations are restricted to the mouse extrastriola, which may be representative of mammalian regular afferent zones: extrastriolar in otolith organs, peripheral in canal organs. It remains to be seen how K_V_1.8 affects properties specific to striolar afferents, such as synchrony and irregularity of firing, which relate to striolar afferents’ enhanced capacity for encoding high-frequency signals (Cullen, 2019).

### K_V_1.8 and vestibulomotor processing

The lack of detectable deficits on the general motor tasks and detectable brain K_V_1.8 expression together support an inner ear origin for the behavioral deficits, involving altered HC physiology and HC-to-afferent transmission. Because K_V_1.8 underlies g_K,L_ in type I HCs and g_A_ and part of g_DR_ in type II HCs, the knockout reveals the combined effect of removing K_V_1.8 across all vestibular HCs. There are also K_V_1.8 channels in cochlear HCs (Dierich et al., 2020), but these are unlikely to play a role in the behaviors and responses that we tested.

The horizontal angular VOR is principally driven by regular afferents from the peripheral zone of the lateral canals, which resembles the otolith extrastriolar zone in physiology and encoding strategy (Schneider et al., 2015). In control mice, VOR responses in darkness (VOR_D_) approached ideal compensatory values at 2 Hz, with gain near 1 and phase near 0. *Kcna10*^-/-^ mice had significantly lower VOR_D_ gain and greater phase deviations at frequencies below 5 Hz. These behavioral deficits occurred despite larger utricular HC receptor potentials and similar low-frequency calyx PSP gain relative to bundle displacement, indicating that larger peripheral responses do not necessarily translate into more effective gain stabilization.

On the balance beam, *Kcna10*^-/-^ mice had more head motion power above 10 Hz, which may reflect impaired head stabilization accompanying their poor balance performance. *Kcna10*^-/-^ mice also developed a compensatory balance strategy, tail wrapping, that has been observed in semiarboreal deer mice while climbing thin branches (Horner, 1954) but has not been previously reported in vestibular mouse mutants. Balance beam deficits have been detected in other vestibular mutants: *Vglut3*^-/-^ mice, which lack quantal glutamatergic transmission (Regalado Núñez et al., 2024; Mukhopadhyay et al., 2026), and *Cyp26b1^-/-^* mice, where the striola has been replaced by extrastriola (Ono et al., 2020).

*Kcna10*^-/-^ mice had abnormal posture in the water, where the absence of stable contact with the ground reduces somatosensory cues and mechanical support for postural control. The aforementioned *Vglut3*^-/-^ and *Cyp26b1^-/-^* mice both swim normally, suggesting that quantal transmission and normal striolar organization are not essential for the swimming performance assessed in those studies. In *Gpr156*^-/-^ mice, the main inner ear effect is the loss of the hair bundle polarity reversal specific to the LES zone. These mice cross beams normally but swim poorly (Ono et al., 2024; Jarysta et al., 2026), perhaps reflecting abnormal head tilt as a result of effects on the LES-to-cerebellum pathway. Among these studies, the contrasting swimming phenotypes are consistent with a contribution of extrastriolar signaling to postural control in water. This is not unexpected as regular (extrastriolar) afferents are associated with signaling head tilt relative to gravity (Goldberg et al., 2012). Righting reflexes appeared qualitatively normal in *Kcna10*^-/-^ pups (P5-P13), however, suggesting some preservation of gravity sensing.

In the water, *Kcna10*^-/-^ mice moved their heads more than control littermates. This increased head motion may reflect impaired stabilization resulting from slower or less reliable vestibular feedback about self-motion. Conversely, in the open field arena, *Kcna10*^-/-^ mice moved their heads *less,* reminiscent of how some patients with vestibular inner ear pathologies tend to keep their heads still (Millar et al., 2023). That *Kcna10*^-/-^ mice also reared less frequently may indicate a general avoidance of unstable postures.

Our results together demonstrate that the K_V_1.8 subunit of mouse utricular HCs enhances the speed and fidelity of vestibular afferent signals about self-motion by making essential contributions to the major outward K_V_ currents in both type I and II HCs. K_V_1.8 deletion substantially reduces the total outwardly rectifying current in both HC types, affecting the receptor potential by reducing bandwidth at its high-frequency end and shifting its operating range. These changes are accompanied at the type I-calyx synapse by reduced synaptic gain (PSP per unit receptor potential) and altered timing (phase) of NQ responses. Vestibulomotor performance is impaired in numerous ways that produce either reduced or less controlled mobility of the head and/or body. The *Kcna10^-/-^* mouse thus provides a molecular tool for linking vestibular hair cell membrane properties to the precision and robustness of gaze and postural control, and for analyzing the unusual nonquantal transmission at the type I HC-calyx synapse.

## Acknowledgements

This study was supported by NIH grant R01 DC012347 (RAE), NSF Graduate Research Fellowship (HRM), UChicago Quad Summer Undergraduate Research Award (ES), 1U01-NS111695 (KEC), and NIH K12-GM123914 (BMV). We thank the Neuroscience Behavioral Core at University of Chicago, particularly Xiaoxi Zhuang, Paschalis Kratsios, and Chris Gomez for use of various facilities. We thank Chenhao Bao and Dale Roberts at Johns Hopkins University for help with the statistical analysis of head motions. We thank Sherri Jones and Tom Friedman for generously donating animals from the *Kcna10*^-/-^ mice introduced in Lee et al. (2013). We thank Zheng-yi Chen for pointing out to us the high expression of *Kcna10* in mouse vestibular hair cell RNAseq data (Scheffer et al., 2015). We thank Radha Kalluri and members of her laboratory, as well as Aravind Chenrayan-Govindaraju and Rob Raphael for insightful feedback on an earlier version of this manuscript.

## Author Contributions

HRM and RAE designed electrophysiological experiments on hair cells and calyces and analyzed data; HRM executed electrophysiological experiments; OLR designed and executed nonquantal transmission experiments; HRM, DS, and ES designed, executed, and analyzed behavioral experiments; BMV designed, executed, and analyzed instrumented behavioral assessments; MG designed, executed, and analyzed VOR; HRM, RAE, and BMV wrote the paper; KEC designed instrumented behavioral and VOR experiments, advised on the analysis, and edited the manuscript. All authors edited the manuscript.

## Abbreviations

*f*_LP_: lowpass corner frequency
g_K,L_: low-voltage-activated K^+^ conductance in type I HCs
g_A_: A-type (inactivating) K_V_ conductance in type II HCs
g_DR_: delayed rectifier K_V_ conductance in type II HCs
HC: hair cell
I_MET_: mechanotransducer current
NQ: nonquantal
VOR: vestibulo-ocular reflex
V_RecP_: receptor potential

## SUPPLEMENTAL ITEMS

**Supplemental Figure 1.**
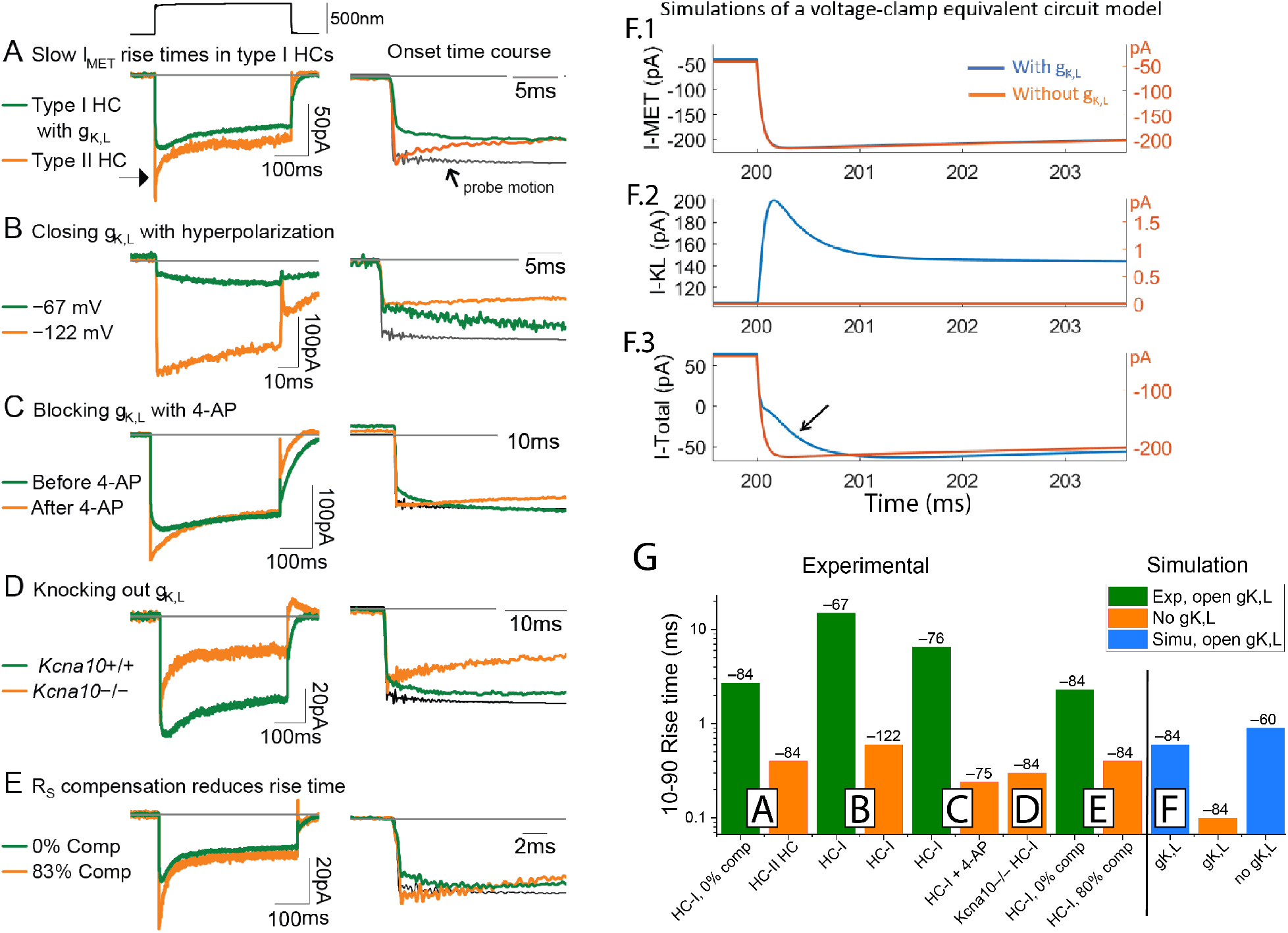
g_K,L_ can introduce creep into voltage-clamp recordings of MET currents. (**A**) MET currents with slow rise times sometimes appeared in 35% (5/14) type I HCs with g_K,L_ but never in type II hair cells (0/9). MET currents had a 10-90% rise time of 2.7 ms in a type I HC and 0.4 ms in a type II HC. Holding potential –84 mV. *Right column*, normalized responses overlaid to show time course at the stimulus onset. (**B**) MET currents’ rise times were reduced by closing g_K,L_ channels by hyperpolarizing below their activation threshold. MET current rise times: at –67 mV, 14.9 ms; at –122 mV, 0.6 ms. (**C**) MET currents’ rise times were reduced by inhibiting g_K,L_ channels with extracellular 5 mM 4-AP. MET currents’ rise times: 6.5 ms before 4-AP; 0.24 ms after 4-AP. Holding potential –74 mV. (**D**) MET currents rise times were smaller in a *Kcna10*^-/-^ type I HC lacking g_K,L_ (0.3 ms) than in a *Kcna10*^+/–^ type I hair cell (2.7 ms). Holding potential –84 mV. (**E**) Series resistance (R_S_) compensation corrected slow rise times. A type I HC (P23 LES, 6pF) with R_S_ = 10 MΩ. MET currents’ rise times: at 0% R_S_ compensation, 2.3 ms; at 80% R_S_ compensation, 0.4 ms. Holding potential –84 mV. (**F**) Simulations of a voltage-clamp equivalent circuit that includes g_MET_, g_K,L_, R_S_, C_m_, and time constants of clamp. MET current (**F.1**) and current through g_K,L_ (**F.2**) are summed (**F.3**) to show current recorded during a whole-cell patch clamp experiment. *Arrow,* creep: the slower, two-phase onset of total current in the presence of g_K,L_. Total current rise times: with g_K,L_, 0.6 ms; without g_K,L._, 0.1 ms. Model parameters in <u>Table S5</u>. (**G**) 10-90% rise times of experimental and simulated MET current. *Labels above each bar*: holding potential (mV). *Boxed letters*: panel of this figure that these data refer to.

**Supplemental Figure 2.**
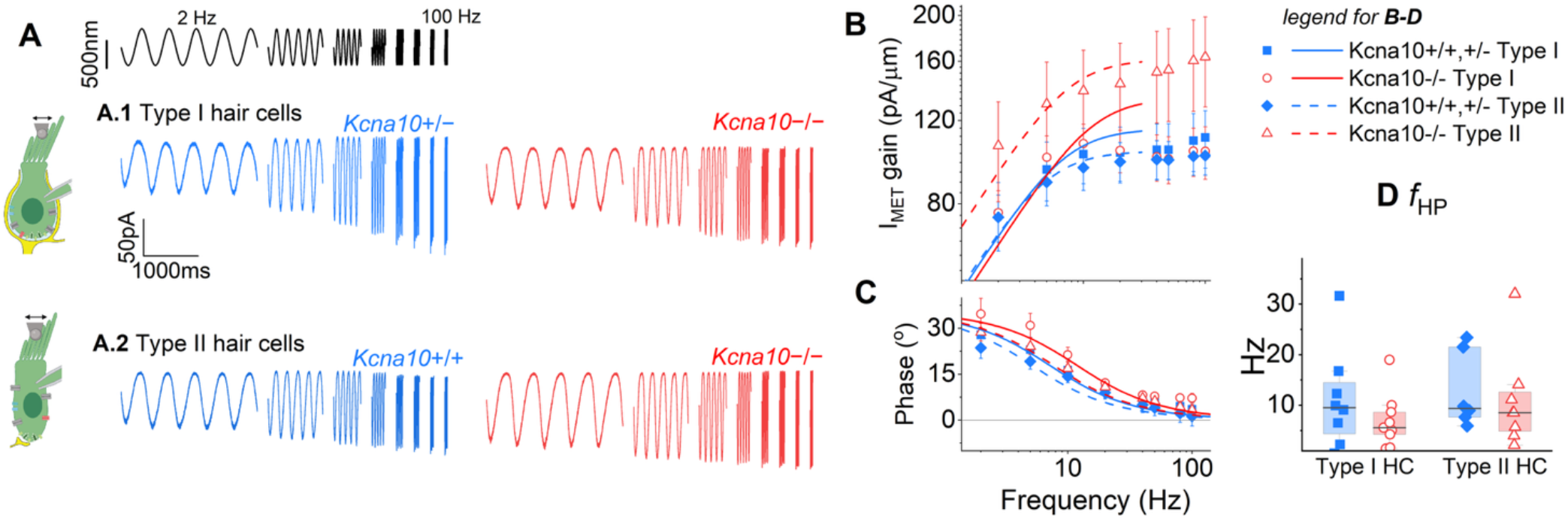
K*c*na10 deletion does not affect frequency tuning of MET currents. (**A**) Exemplar MET currents from type I HCs (***A.1***, *left*, P17 LES, V_hold_ −74 mV; *right*, P45 MES, V_hold_ −84 mV) and type II HCs (***A.2***, *left*, P22 MES, V_hold_ −74 mV; *right*, P49 MES, V_hold_ −74 mV). Each trace averages 4-14 presentations. Bundle displacement at top and scale bar in ***A.1*** apply to whole figure. (**B**) Peak-to-peak (AC) gain fit to Eq. 4a. Type I HCs, mixed ANOVA: Genotype, p=0.98; Frequency, p=8E-4; Genotype-Frequency Interaction, p=0.47. Type II HCs, mixed ANOVA: Genotype, p=0.13; Frequency, p=4E-6; Genotype-Frequency Interaction, p=0.03*. (**C**) Phase fit to Eq. 4b. Type I HCs, mixed ANOVA: Genotype, p = 0.3; Frequency, p = 8E-8; Genotype-Frequency Interaction, p = 0.4. Type II HCs, mixed ANOVA: Genotype, p = 0.37; Frequency, p = 2E-10; Genotype-Frequency Interaction, p = 0.6. (**D**) Highpass corner frequencies. *Line*, medians. See Table 4 for statistics.

---

**Supplemental Table 1.** VOR statistical tests at each frequency.

|  | Group<br>ANOVA<br>(n mice) | Group<br>ANOVA<br>(n mice) | 0.4 Hz | 0.6 Hz | 0.8 Hz | 1 Hz | 2 Hz | 5 Hz | 10 Hz | 15 Hz | 20 Hz |
|---|---|---|---|---|---|---|---|---|---|---|---|
| VORd | WT (4)<br>*vs*<br>HET (10) | Ctrl (14)<br>*vs*<br>Null (9) |  |  |  |  |  |  |  |  |  |
| Gain | p = 0.87 | p = 2E-4 | 9E-6 | 1.5E-5 | 2E-4 | 2E-4 | 0.012 | 0.04 | 0.8 | 0.58 | 0.3 |
| Phase | p = 0.91 | p = 7E-7 | 5E-4 | 1.4E-5 | 1.8E-5 | 2.9E-5 | 0.04 | 0.98 | 0.15 | 0.41 | 0.3 |
| VORl | WT (4)<br>*vs*<br>HET (8) | Ctrl (12)<br>*vs*<br>Null (9) |  |  |  |  |  |  |  |  |  |
| Gain | p = 0.31 | p = 2E-7 | 0.05 | 0.035 | 0.033 | 0.043 | 0.036 | 0.51 | 0.95 | 0.47 | 0.09 |
| Phase | p = 0.99 | p = 0.02 | 0.5 | 0.16 | 0.21 | 0.12 | 0.022 | 0.16 | 0.72 | 0.45 | 0.12 |

**Supplemental Table 2.**
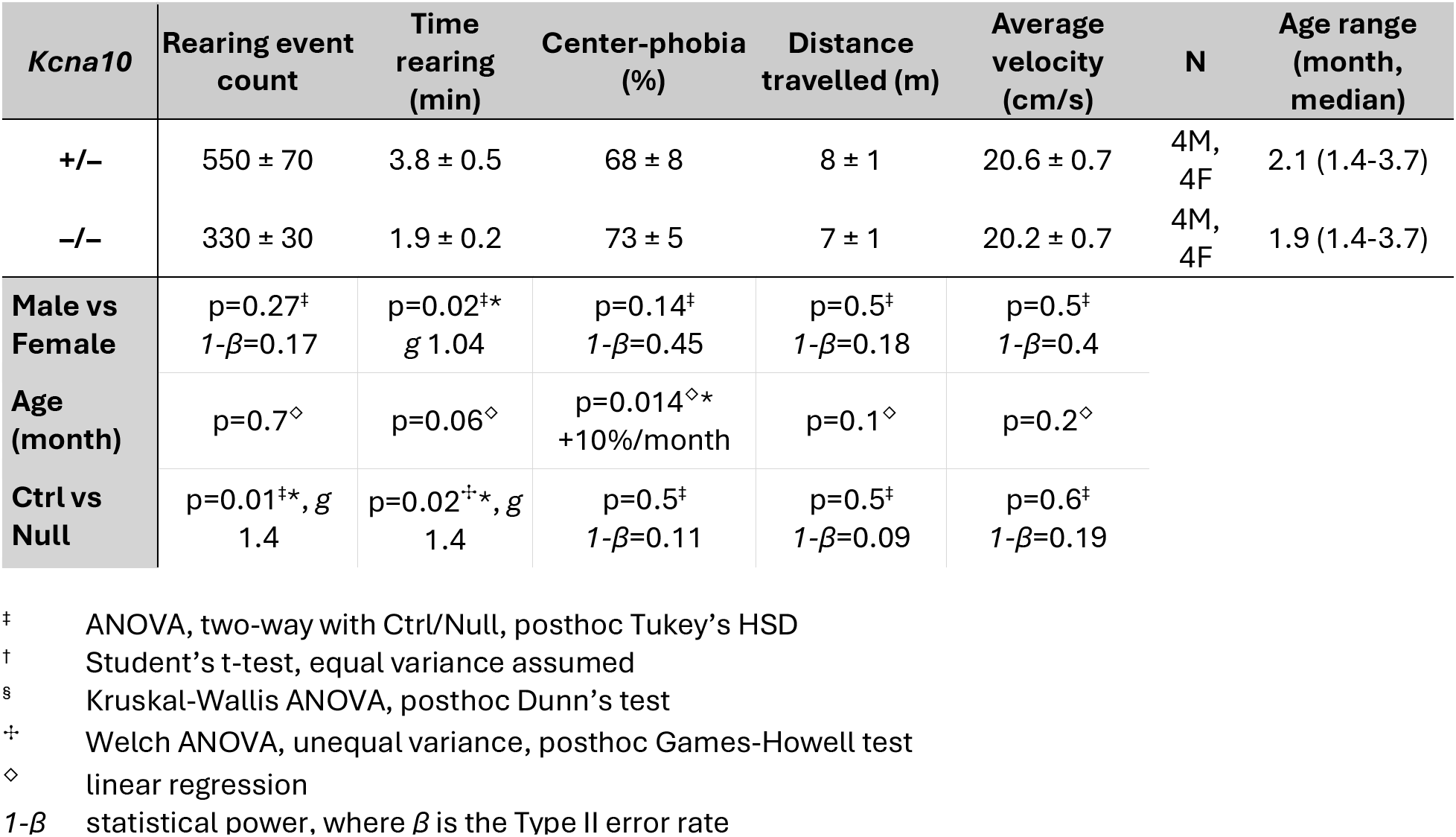
Exploration of an open field arena for 1 hour. Mean ± SEM (N is number of mice). Age is median months (range)

| ***Kcna10*** | **Rearing event count** | **Time rearing (min)** | **Center-phobia (%)** | **Distance travelled (m)** | **Average velocity (cm/s)** | **N** | **Age range (month, median)** |
|---|---|---|---|---|---|---|---|
| **+/–** | 550 ± 70 | 3.8 ± 0.5 | 68 ± 8 | 8 ± 1 | 20.6 ± 0.7 | 4M, 4F | 2.1 (1.4-3.7) |
| **–/–** | 330 ± 30 | 1.9 ± 0.2 | 73 ± 5 | 7 ± 1 | 20.2 ± 0.7 | 4M, 4F | 1.9 (1.4-3.7) |
| **Male vs Female** | p=0.27<sup>‡</sup><br>*1-β*=0.17 | p=0.02<sup>‡</sup>\*<br>*g* 1.04 | p=0.14<sup>‡</sup><br>*1-β*=0.45 | p=0.5<sup>‡</sup><br>*1-β*=0.18 | p=0.5<sup>‡</sup><br>*1-β*=0.4 |  |  |
| **Age (month)** | p=0.7<sup>◇</sup> | p=0.06<sup>◇</sup> | p=0.014<sup>◇</sup>*<br>+10%/month | p=0.1<sup>◇</sup> | p=0.2<sup>◇</sup> |  |  |
| **Ctrl vs Null** | p=0.01<sup>‡</sup>\*, *g* 1.4 | p=0.02<sup>✢</sup>\*, *g* 1.4 | p=0.5<sup>‡</sup><br>*1-β*=0.11 | p=0.5<sup>‡</sup><br>*1-β*=0.09 | p=0.6<sup>‡</sup><br>*1-β*=0.19 |  |  |

<sup>‡</sup> ANOVA, two-way with Ctrl/Null, posthoc Tukey’s HSD
<sup>†</sup> Student’s t-test, equal variance assumed
<sup>§</sup> Kruskal-Wallis ANOVA, posthoc Dunn’s test
<sup>✢</sup> Welch ANOVA, unequal variance, posthoc Games-Howell test
<sup>◇</sup> linear regression
*1-β* statistical power, where *β* is the Type II error rate

**Supplemental Table 3.**
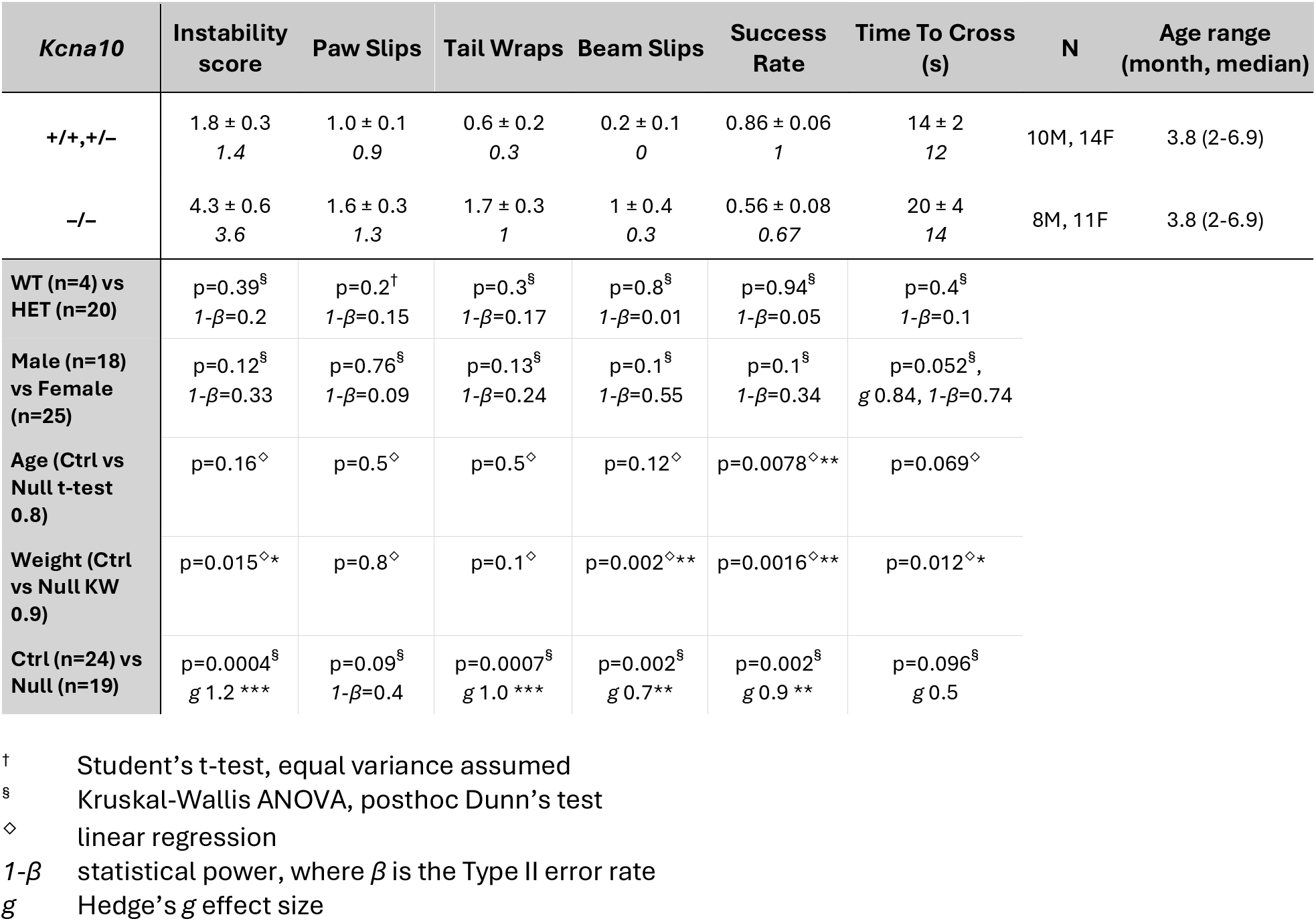
Ǫuantitative descriptions of crossing a narrow balance beam in K*cna10^+/+,^*^+/–^ and *Kcna10*^-/-^ mice. Mean ± SEM (N is number of mice). Age is median months (range). Median (Med) is provided for non-normally distribut

| *Kcna10* | **Instability score** | **Paw Slips** | **Tail Wraps** | **Beam Slips** | **Success Rate** | **Time To Cross (s)** | **N** | **Age range (month, median)** |
|---|---|---|---|---|---|---|---|---|
| **+/+,+/–** | 1.8 ± 0.3<br>*1.4* | 1.0 ± 0.1<br>*0.9* | 0.6 ± 0.2<br>*0.3* | 0.2 ± 0.1<br>*0* | 0.86 ± 0.06<br>*1* | 14 ± 2<br>*12* | 10M, 14F | 3.8 (2-6.9) |
| **–/–** | 4.3 ± 0.6<br>*3.6* | 1.6 ± 0.3<br>*1.3* | 1.7 ± 0.3<br>*1* | 1 ± 0.4<br>*0.3* | 0.56 ± 0.08<br>*0.67* | 20 ± 4<br>*14* | 8M, 11F | 3.8 (2-6.9) |
| **WT (n=4) vs HET (n=20)** | p=0.39<sup>§</sup><br>*1-β*=0.2 | p=0.2<sup>†</sup><br>*1-β*=0.15 | p=0.3<sup>§</sup><br>*1-β*=0.17 | p=0.8<sup>§</sup><br>*1-β*=0.01 | p=0.94<sup>§</sup><br>*1-β*=0.05 | p=0.4<sup>§</sup><br>*1-β*=0.1 |  |  |
| **Male (n=18) vs Female (n=25)** | p=0.12<sup>§</sup><br>*1-β*=0.33 | p=0.76<sup>§</sup><br>*1-β*=0.09 | p=0.13<sup>§</sup><br>*1-β*=0.24 | p=0.1<sup>§</sup><br>*1-β*=0.55 | p=0.1<sup>§</sup><br>*1-β*=0.34 | p=0.052<sup>§</sup>,<br>*g* 0.84, *1-β*=0.74 |  |  |
| **Age (Ctrl vs Null t-test 0.8)** | p=0.16<sup>◇</sup> | p=0.5<sup>◇</sup> | p=0.5<sup>◇</sup> | p=0.12<sup>◇</sup> | p=0.0078<sup>◇</sup>** | p=0.069<sup>◇</sup> |  |  |
| **Weight (Ctrl vs Null KW 0.9)** | p=0.015<sup>◇</sup>* | p=0.8<sup>◇</sup> | p=0.1<sup>◇</sup> | p=0.002<sup>◇</sup>** | p=0.0016<sup>◇</sup>** | p=0.012<sup>◇</sup>* |  |  |
| **Ctrl (n=24) vs Null (n=19)** | p=0.0004<sup>§</sup><br>*g* 1.2 \*\*\* | p=0.09<sup>§</sup><br>*1-β*=0.4 | p=0.0007<sup>§</sup><br>*g* 1.0 \*\*\* | p=0.002<sup>§</sup><br>*g* 0.7\*\* | p=0.002<sup>§</sup><br>*g* 0.9 \*\* | p=0.096<sup>§</sup><br>*g* 0.5 |  |  |

<sup>†</sup> Student’s t-test, equal variance assumed
<sup>§</sup> Kruskal-Wallis ANOVA, posthoc Dunn’s test
◇ linear regression
*1-β* statistical power, where *β* is the Type II error rate
*g* Hedge’s *g* effect size

**Supplemental Table 4.**
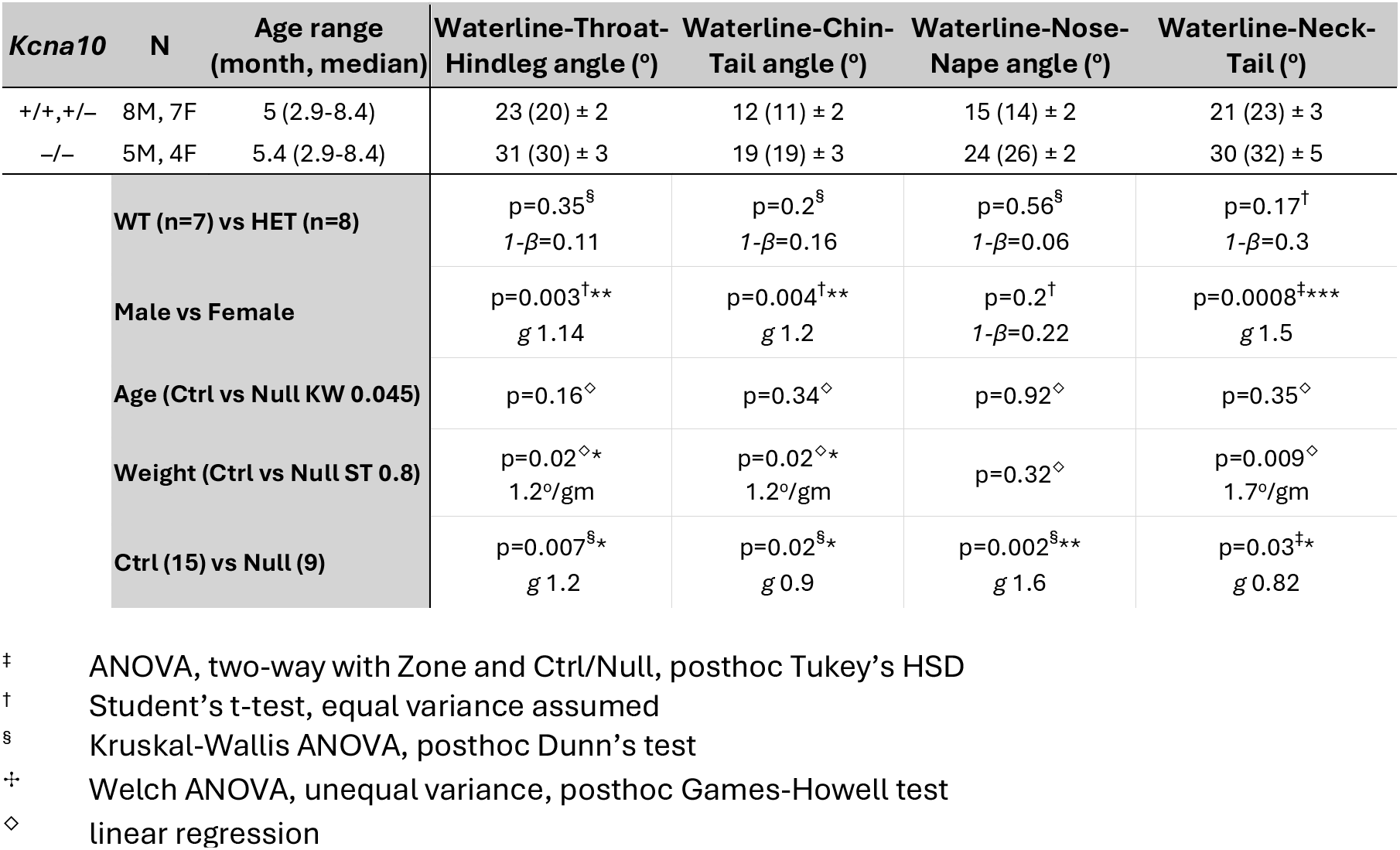
Swim body posture differed between Kcna10+/– and Kcna10-/- mice. Mean (median) ± SEM.

| *Kcna10* | N | Age range<br>(month, median) | Waterline-Throat-Hindleg angle (º) | Waterline-Chin-Tail angle (º) | Waterline-Nose-Nape angle (º) | Waterline-Neck-Tail (º) |
|---|---|---|---|---|---|---|
| +/+,+/– | 8M, 7F | 5 (2.9-8.4) | 23 (20) ± 2 | 12 (11) ± 2 | 15 (14) ± 2 | 21 (23) ± 3 |
| –/– | 5M, 4F | 5.4 (2.9-8.4) | 31 (30) ± 3 | 19 (19) ± 3 | 24 (26) ± 2 | 30 (32) ± 5 |
|  | **WT (n=7) vs HET (n=8)** |  | p=0.35<sup>§</sup><br>*1-β*=0.11 | p=0.2<sup>§</sup><br>*1-β*=0.16 | p=0.56<sup>§</sup><br>*1-β*=0.06 | p=0.17<sup>†</sup><br>*1-β*=0.3 |
|  | **Male vs Female** |  | p=0.003<sup>†</sup>\*\*<br>*g* 1.14 | p=0.004<sup>†</sup>\*\*<br>*g* 1.2 | p=0.2<sup>†</sup><br>*1-β*=0.22 | p=0.0008<sup>‡</sup>\*\*\*<br>*g* 1.5 |
|  | **Age (Ctrl vs Null KW 0.045)** |  | p=0.16<sup>◇</sup> | p=0.34<sup>◇</sup> | p=0.92<sup>◇</sup> | p=0.35<sup>◇</sup> |
|  | **Weight (Ctrl vs Null ST 0.8)** |  | p=0.02<sup>◇</sup>*<br>1.2º/gm | p=0.02<sup>◇</sup>*<br>1.2º/gm | p=0.32<sup>◇</sup> | p=0.009<sup>◇</sup><br>1.7º/gm |
|  | **Ctrl (15) vs Null (9)** |  | p=0.007<sup>§</sup>\*<br>*g* 1.2 | p=0.02<sup>§</sup>\*<br>*g* 0.9 | p=0.002<sup>§</sup>\*\*<br>*g* 1.6 | p=0.03<sup>‡</sup>\*<br>*g* 0.82 |

<sup>‡</sup> ANOVA, two-way with Zone and Ctrl/Null, posthoc Tukey’s HSD
<sup>†</sup> Student’s t-test, equal variance assumed
<sup>§</sup> Kruskal-Wallis ANOVA, posthoc Dunn’s test
<sup>✢</sup> Welch ANOVA, unequal variance, posthoc Games-Howell test
<sup>◇</sup> linear regression

**Supplemental Table 5.** Model parameters for the voltage-equivalent circuit used for simulations of current through MET and g_K,L_ channels in Fig. S1. Model adapted from Lei et al., 2020.

| Symbol | Value | Description |
|---|---|---|
| **Capacitance** |  |  |
| *C*<sub>f</sub> | 0.3 pF | Feedback shunt capacitance(Lei et al., 2020) |
| *C*<sub>m</sub> | 5 pF | Membrane capacitance |
| *C*<sub>p</sub> | 3-5 pF |  |
| **Current** |  |  |
| *I*<sub>in</sub> |  | Voltage-clamp current |
| *I*<sub>ion</sub> |  | Whole-cell ion channel current |
| *I*<sub>leak</sub> |  | Leakage current through imperfect seal |
| *I*<sub>m</sub> |  | Membrane current |
| *I*<sub>out</sub> |  | Recorded current |
| **Resistance** |  |  |
| *R*<sub>f</sub> | 500 MΩ | Feedback resistance (current gain on HEKA EPC 10) |
| *R*<sub>S</sub> | 10 MΩ | Series resistance between pipette electrode and cell |
| *α* | 0-0.8 | Fraction of R<sub>S</sub> compensation |
| *R*<sub>seal</sub> | 0.5-10 GΩ | Seal resistance of the pipette tip |
| **Time constant** |  |  |
| *τ*<sub>a</sub> | 50 μs | Membrane access time constant, *τ*<sub>a</sub> = *R*<sub>s</sub>*C*<sub>m</sub> |
| *τ*<sub>sum</sub> | 40 μs | Response time of the summing amplifier(Lei et al., 2020) |
| *τ*<sub>clamp</sub> | 10 μs | Voltage-clamp time constant, *τ*<sub>clamp</sub> = (1 – *α*)\* *τ*<sub>a</sub> |
| *τ*<sub>z</sub> | 150 μs | Transconductor time constant, *τ*<sub>z</sub> = *R*<sub>f</sub>*C*<sub>f</sub> |
| **Voltage** |  |  |
| *V*<sub>cmd</sub> |  | Command voltage, follows the voltage-clamp protocol |
| *V*<sub>clamp</sub> |  | Clamp voltage |
| *V*<sub>m</sub> |  | Membrane potential |
| *V*<sub>off</sub> |  | Offset voltage: amplifier offsets, electrode offsets, junction potentials |
| *V*<sub>out</sub> |  | Recorded voltage, *V*<sub>out</sub> = *I*<sub>out</sub>*R*<sub>f</sub> |
| *V*<sub>p</sub> |  | Pipette potential |
| **Conductance: g<sub>K,L</sub>** |  |  |
| *g*<sub>max</sub> | 300 nS | Maximal conductance (Martin et al., 2024) |
| *V*<sub>half</sub> | –84 mV | Midpoint voltage activation |
| *dV* | 5 mV | Slope factor of voltage-dependence |
| *τ*<sub>act</sub> | 300 ms | Time constant of activation |
| *E*<sub>K</sub> | –86 mV | K<sup>+</sup> reversal potential |
| **Conductance: g<sub>MET</sub>** |  |  |
| *g*<sub>max</sub> | 3 nS | Maximal conductance |
| *X*<sub>1/2</sub> | 0.3 μm | Midpoint displacement activation |
| *dX* | 0.2 μm | Slope factor of displacement sensitivity |
| *τ*<sub>Act</sub> | 50 μs | Activation time constant |
| *τ*<sub>Adapt</sub> | 10 ms | Adaptation time constant |
| *E*<sub>met</sub> | 2.6 mV | I<sub>MET</sub> reversal potential (Kros et al., 1992) |
| **Leak through seal** |  |  |
| *E*<sub>Leak</sub> |  | V<sub>m</sub> |
| *g*<sub>Leak</sub> | 2 nS | 1/*R*<sub>Seal</sub>, (Lei et al., 2020) |

**Supplemental Table 6.** Distribution of sex among electrophysiology recordings of individual cells.

|  |  | **Total mice** | **Female** | **Male** | **Unknown Sex** | **Place in manuscript with data** |
|---|---|---|---|---|---|---|
| **Type I** | Control | 8 | 2 | 6 | 0 | Figures 1-2; Tables 1-2,4 |
|  | Null | 8 | 7 | 1 | 0 |  |
| **Type II** | Control | 9 | 3 | 3 | 3 | Figures 1-2; Tables 1-2,4 |
|  | Null | 8 | 5 | 3 | 0 |  |
| **Calyx** – voltage-gated | Control | 34 | 9 | 19 | 6 | Figure 3; Table 3 |
|  | Null | 22 | 6 | 15 | 1 |  |
| **Calyx** – synaptic transmission | Control | 5 | 0 | 1 | 5 | Figure 4; Table 4 |
|  | Null | 6 | 1 | 3 | 2 |  |

**Supplemental Table 7.** Type I and type II datasets were combined to increase the power of detecting differences based on type, sex, and zone. We included interaction terms in the model only if we expected them based on findings in the main analysis. The multi-factor ANOVA used the type II sum-of-squares method because it is more robust to unbalanced cohorts, which was especially true for sex and zone. If a main effect was found in the model, then posthoc pairwise p-values are reported below the table. If the pairwise comparison involved a non-normally distributed group, then a Kruskal-Wallis ANOVA (KWA) was used. If the pairwise comparison involved unequal variance, then a Welch ANOVA was used.

| **Parameter** | **Control *vs* Null p-value** | **Type I *vs* type II p-value** | **Male *vs* Female p-value** | **MES *vs* LES p-value** | **Interactions were included in model if driven by a hypothesis based on the main analysis** |
|---|---|---|---|---|---|
| G<sub>met</sub> (nS) | 0.06 | 0.4 | 0.9 | 0.6 |  |
| I<sub>MET</sub> X<sub>half</sub> (um) | 0.3 | 0.07 | 0.7 | 0.2 |  |
| I<sub>MET</sub> slope factor (um) | 0.3 | 0.5 | 0.2 | 0.9 |  |
| P<sub>0</sub> (%) | 0.63 | 0.01*<sup>a</sup> | 0.3 | 0.02*<sup>b</sup> | Age*Type interaction included: p=0.2 |
| V<sub>RecP</sub> X<sub>half</sub> (um) | 0.03*<sup>c</sup> | 0.07 | 0.96 | 0.4 | Type*Kcna10 interaction included: p=0.03*<sup>c</sup> |
| V<sub>RecP</sub> slope factor (um) | 0.6 | 0.4 | 0.7 | 0.3 |  |
| V<sub>RecP</sub> Tau (ms) | 0.001**<sup>d</sup> | 0.047*<sup>e</sup> | 0.6 | 0.8 | Type*Kcna10 interaction included: p=0.7 |

<sup>a</sup> Type I, 10.2 ± 0.4 % (n=16) *vs* Type II, 20.8 ± 0.9 % (n=15); Hedge’s *g* = 1; Welch ANOVA p = 0.01
<sup>b</sup> MES, 22 ± 3 % (n=12) *vs* LES, 11 ± 2 % (n=19); Hedge’s *g* = 1; KWA p = 0.009
<sup>c</sup> Given a significant interaction, we report the posthoc pairwise p-values adjusted with Tukey for multiple comparisons
Control type I vs null type I, p=0.0006; control type I vs control type II, p=0.003; control type I vs null type II, p=0.002. Thus, the effect is based on the presence or absence of g<sub>K,L</sub>
<sup>d</sup> Control, 2.1 ± 0.6 ms (n=14, median 1.2) *vs* Null, 8 ± 1 ms (n=14); Hedge’s *g* 1.5; KWA p=0.0004
<sup>e</sup> Type I, 3 ± 1 ms (n=14, median 1.4) *vs* Type II, 6 ± 1 ms (n=14); Hedge’s *g* = 0.7; KWA p = 0.048

